# Advancing image-based meta-analysis for fMRI: A framework for leveraging NeuroVault data

**DOI:** 10.1101/2025.03.06.641922

**Authors:** Julio A. Peraza, James D. Kent, Ross W. Blair, Jean-Baptiste Poline, Thomas E. Nichols, Alejandro de la Vega, Angela R. Laird

## Abstract

Image-based meta-analysis (IBMA) is a powerful method for synthesizing results from various fMRI studies. However, challenges related to data accessibility and the lack of available tools and methods have limited its widespread use. This study examined the current state of the NeuroVault repository and developed a comprehensive framework for selecting and analyzing neuroimaging statistical maps within it. By systematically assessing the quality of NeuroVault’s data and implementing novel selection and meta-analysis techniques, we demonstrated the repository’s potential for IBMA. We created a multi-stage selection framework that includes preliminary, heuristic, and manual image selection methods. We conducted meta-analyses for three distinct domains: working memory, motor, and emotion processing. The results from the three manual IBMA approaches closely resembled reference maps from the Human Connectome Project. Importantly, we found that while manual selection provides the most precise results, heuristic methods can serve as robust alternatives, especially for domains that include a heterogeneous set of fMRI tasks and contrasts, such as emotion processing. Additionally, we evaluated five different meta-analytic estimator methods to assess their effectiveness in handling spurious images. For domains characterized by heterogeneous tasks, employing a robust estimator (e.g., median) is essential. This study is the first to present a systematic approach for implementing IBMA using the NeuroVault repository. We introduce an accessible and reproducible methodology that allows researchers to make the most of NeuroVault’s extensive neuroimaging resources, potentially fostering greater interest in data sharing and future meta-analyses utilizing NeuroVault data.

## Introduction

Despite the increasing adoption of NeuroVault as a community-based neuroimaging repository, downloading usable data for image-based meta-analysis (IBMA) still presents multiple challenges. For example, a substantial portion of NeuroVault collections are annotated incorrectly or lack a link to a valid publication; some images are duplicates, while others correspond to non-statistical imaging modalities (Menuet et al., 2022). Here, we explored the current state of NeuroVault, identified different challenges in the archived data, and proposed an automatic heuristic selection framework for curating images for IBMA.

IBMA, considered the gold standard of neuroimaging meta-analysis (Salimi-Khorshidi et al., 2009; Salo et al., 2023), consists of aggregating results from group-level, whole-brain statistical maps from individually conducted functional magnetic resonance imaging (fMRI) studies. IBMA outperforms other popular meta-analysis methods, such as coordinate-based meta-analysis (CBMA) (Fox et al., 1999, 1998, 1997; Nielsen and Hansen, 2002). For instance, CBMA works with activation foci producing information loss and it relies on kernel-based methods to infer the model activation maps from peak coordinates, requiring spatial assumptions that fail to reproduce the actual statistical map (Wager et al., 2007). Although a combination of kernel parameters and thresholds for CBMA could be chosen to maximize the similarity with IBMA maps, such optimal settings depend on the dataset and specific considerations from individual studies (e.g., statistical thresholds) and thus are not generalizable across meta-analyses (Salimi-Khorshidi et al., 2009). Further, considering how underpowered most neuroimaging studies are, brain activations that fail to pass certain thresholds of significance are generally discarded in a CBMA. In contrast, IBMA methods use whole-brain statistics; thus, all existing voxel-wise statistical methods are available to analyze subject-level data within studies (Lazar et al., 2002). IBMA is known to produce richer and more detailed results, with additional brain structures that are often absent from CBMA results. IBMA also has greater power; thus, one could potentially achieve similar or even better results with a small fraction of studies generally required in CBMA. In addition, when both the parameters and variance estimates are available, hierarchical mixed effect models can be used to account for both within- and between-study variance (Salimi-Khorshidi et al., 2009). Despite the clear superiority of IBMA, CMBA remains the most popular approach among the neuroimaging research community.

In the last two decades, only a few IBMAs have been conducted. Some researchers have leveraged IBMA methodologies for combining data from multiple sites from big data consortiums, such as ABIDE (Ma et al., 2023) and the Brainnetome Project for Schizophrenia (Cui et al., 2022). Other studies have followed a more traditional approach to meta-analysis by identifying relevant studies in the literature and then contacting the corresponding author of the selected publications, asking for the group-level unthresholded statistical maps (Fiorito et al., 2023; Hellewell et al., 2020; Lamm et al., 2011; Luijten et al., 2017; Lukow et al., 2021; Schulze et al., 2019, 2016; Witt et al., 2021). Limiting IBMA to neuroimaging consortiums restricts meta-analytic topics to a few domains and scientific questions. Alternatively, traditional meta-analysis represents a more arduous task since contacting several dozen scientists and asking for their group-level unthresholded and normalized statistical maps is time-consuming. Moreover, corresponding authors are not always responsive; the data could be lost, difficult to retrieve, or lack the quality required for a meta-analysis (Fiorito et al., 2023). In comparison, more than 400 CBMA studies have been conducted since 2010, addressing consensus for many domains and scientific questions. In practice, researchers commonly report activation foci of significant findings in neuroimaging studies, making a large portion of the neuroimaging literature accessible for CBMA approaches (Fox et al., 2005; Laird et al., 2011, 2009, 2005). Meanwhile, whole-brain statistical images are rarely shared, thus limiting IBMA to a small number of tasks and mental functions.

The NeuroVault web-based repository was introduced to address image data availability by providing an easy-to-use community platform to share statistical maps (Gorgolewski et al., 2015). Today, NeuroVault’s archives contain a significant volume of collections with more than two hundred thousand brain maps and corresponding metadata, some of which are linked to peer-reviewed articles. However, there is still limited coverage of the neuroimaging literature, as most research labs have not adopted NeuroVault as part of their workflows when submitting an article for publication (Salo et al., 2023). Currently, downloading usable data from NeuroVault comes with multiple challenges. For example, a substantial portion of collections in NeuroVault are wrongly annotated or lack a link to a valid publication; some images are duplicates, and others correspond to non-statistics imaging modalities (Menuet et al., 2022). Overall, the potential number of spurious statistical maps complicates the use of NeuroVault data for IBMA. Although the NeuroVault API is integrated with some neuroimaging software (e.g., Nilearn and NiMARE) (Salo et al., 2023), users without coding experience often struggle to produce efficient analyses from NeuroVault data. Taken together, there is a need for standard guidelines for image selection and proper data cleaning, followed by open-source tools and reproducible methods to facilitate IBMA with NeuroVault.

The overall objective of the current study was to advance research tools for IBMA, including standard image selection framework, image aggregation methods, databases, and guidelines. Initially, we examined the current status of the NeuroVault repository and evaluated its feasibility for IBMA. Then, we developed an image selection framework to identify images suitable for IBMA for a given domain or fMRI tasks. We implemented several combination methods with robust approaches to handle images with extreme values and outliers. The different combinations of image selection and combination methods were assessed against reference images from the task-fMRI group-average effect size maps from the Human Connectome Project (HCP) S1200 data release (Barch et al., 2013; Smith et al., 2013; Uğurbil et al., 2013; Van Essen et al., 2013, 2012). The comparisons between our IBMA results and the reference maps focused on evaluating image similarity and increased estimates in specific brain regions of interest. The entire process, from accessing data in NeuroVault to producing meta-analytic maps, is detailed in an open-access repository to facilitate reproducible and systematic meta-analysis. Collectively, we expect the results of the current work, including our specific guidelines, tools, and methods, to boost interest in the NeuroVault platform and promote IBMA research.

## Results

### Overview

Figure 1 provides an overview of our methodological approach. First, we identified fMRI tasks linked to a specific domain (e.g., working memory) using the established connection between NeuroVault and the Cognitive Atlas knowledge base (Gorgolewski et al., 2015; Poldrack et al., 2011). Then, we downloaded all images linked to selected tasks from the NeuroVault repository. Second, we performed a preliminary image selection leveraging the metadata associated with the images, which were identified as potential candidates for IBMA. We also conducted a data-driven heuristic selection to remove possible outliers from the data. Subsequently, we manually selected relevant images by identifying the analysis contrast in their corresponding article and with the help of the image metadata in NeuroVault. Third, we conducted image-based meta-analyses using standardized effect size maps of the chosen images. In addition to using a baseline meta-analytic estimator (i.e., mean), also referred to in this article as combination or aggregation, we explored four robust combination methods: median, trimmed mean, winsorized mean, and weighted mean. Finally, the meta-analyses with different combinations of parameters (i.e., image selection method and estimator approach) were evaluated against reference images from the task-fMRI group-average effect size maps from the Human Connectome Project (HCP) S1200 data release (Barch et al., 2013; Smith et al., 2013; Uğurbil et al., 2013; Van Essen et al., 2013, 2012).

**Figure 1.**
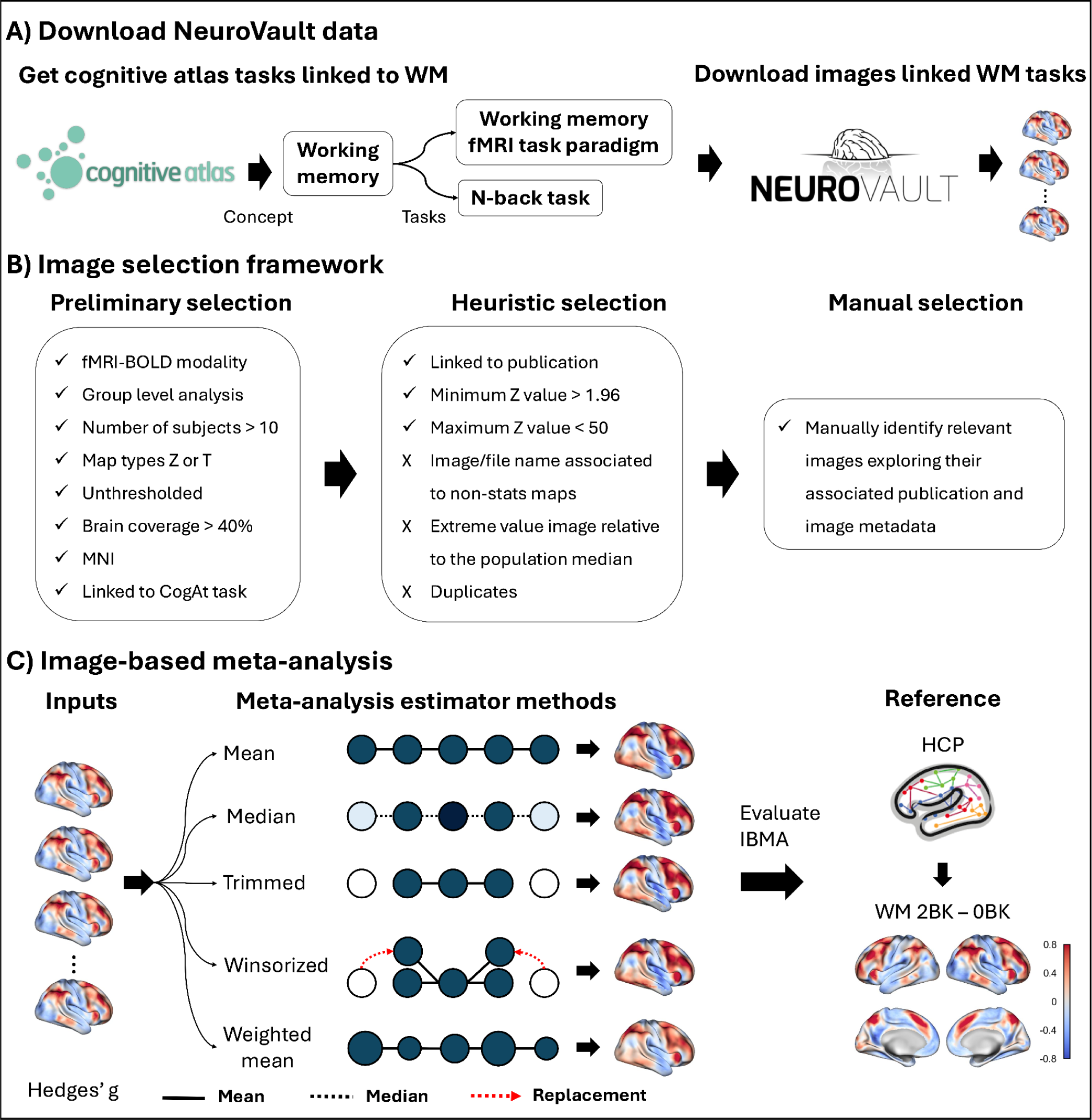
Overview of the general framework for performing image-based meta-analysis with NeuroVault data. **(A)** We identified tasks that measure a given domain of interest using the established connection in the Cognitive Atlas. Then, we downloaded images and metadata associated with the selected tasks from NeuroVault. **(B)** Within the image selection framework, preliminary selection includes images that were potentially suitable for IBMA, such as fMRI-BOLD modality, group-level analysis, sample size larger than ten, z- and t-stats maps, unthresholded, with a brain coverage larger than 40%, and on MNI space. A data-driven heuristic selection was then applied to remove potential outliers from the data. A manual selection process was conducted to find the most relevant images linked to the articles. **(C)** In addition to the baseline estimator (i.e., mean), four different robust methods, such as median, trimmed, winsorized, and weighted mean, were implemented to address extreme values and outliers in the data. The output of the IBMAs was compared with reference maps from the HCP data. The circles represent studies, and the intensity of the color marks influences the total aggregation. Darker colors weigh more in the aggregation, while white circles are discarded studies. The size of the circler, also representing the influence of a study, is different only for the weighted mean. Solid lines represent the mean of the connected circles, while the dotted line denotes the median. For the winsorized mean, a dotted red arrow represents the replacement of extreme values with their nearest neighbors.

### The current state of NeuroVault

Before attempting to run IBMA, we conducted a thorough analysis of the state of NeuroVault as of February 2024. Figure 2A illustrates its evolution from 2013 to 2023. When NeuroVault was launched, it received 11 collections and 63 images in the first year. Since then, the amount of uploaded data has steadily increased, reaching over 238,000 images distributed across 5,756 collections a decade later. Of these collections, 26% are linked to published journal articles, and approximately 66% of images are associated with a publication. The growth of collections follows a similar trend, with a median of 618 new collections created each year. Notably, the most significant increases occurred in 2018 and 2023, with more than 700 collections added each year. The median number of images uploaded annually exceeded 11,000, with the highest uploads occurring in 2018, 2020, and 2021, totaling over 47,000, 54,000, and 58,000 images, respectively. Overall, these trends highlight the rapid growth and increasing adoption of NeuroVault as a comprehensive and widely used repository for sharing neuroimaging statistical images.

**Figure 2.**
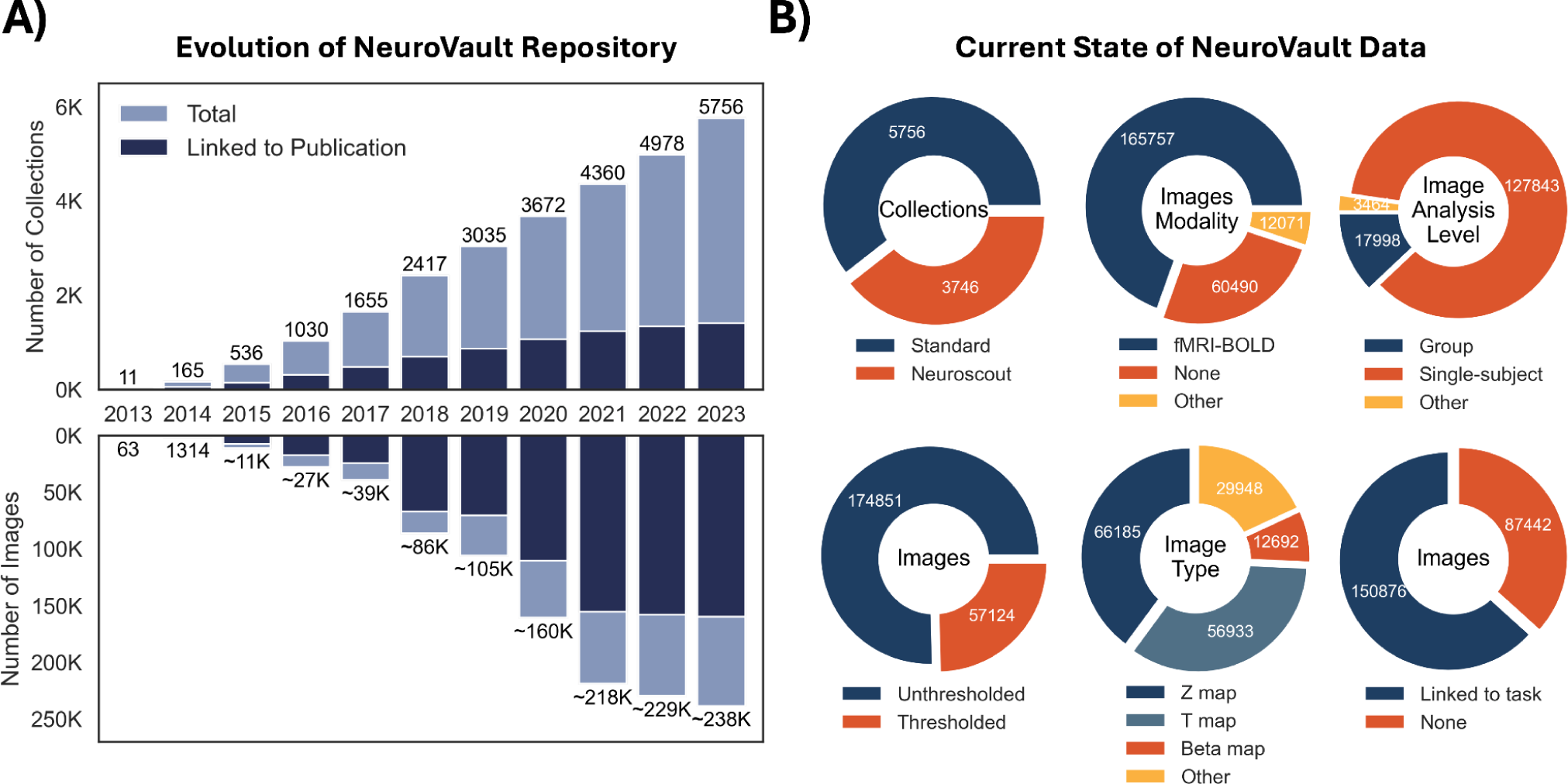
Evolution and current state of NeuroVault data. **(A)** Evolution of the NeuroVault repository from its launch in 2013 until 2023. The top bar plot represents the number of cumulative collections, along with the number of collections from the total that are linked to published scientific articles with a valid DOI. The bottom bar plot presents the cumulative number of images and the subset associated with published articles. **(B) Top Left:** Proportion of standard NeuroVault collections and those created by third-party apps, such as Neuroscout. **Top Middle:** Proportion of different image modalities, “Other” includes diffusion MRI, EEG, MEG, PET FDG, PET other, structural MRI, fMRI-CBF, and fMRI-CBV, while “None” represents missing data or the modality is just not specified in the metadata. **Top Right:** Proportion of the image analysis level. “Other” includes images resulting from a meta-analysis and others. **Bottom Left:** Proportion of unthresholded to thresholded images. **Bottom Middle:** Proportion on images by type, Z, T, Beta, and other maps. “Other” includes F, chi-squared, P, multivariate-beta, ROI/mask, parcellation, anatomical, and variance, amongst other maps. **Bottom Right:** Proportion of images linked to Cognitive Atlas tasks. “None” represents missing data or tasks not specified in the metadata.

Next, we sought to identify which collections could be linked to publications with a valid DOI. Starting with NeuroVault’s 5,756 collections, we found only 896 with a valid DOI link in the DOI field of the collection metadata. Of the remaining 4,985, we found 28 collections with a valid DOI link provided in the collection description field. We then searched PubMed for articles that matched the collection’s title and found 354 additional collections that could be linked to published articles. Finally, we conducted an extensive search using Pubget, an open-source Python tool for collecting data for biomedical text mining. We performed a Pubget query and retrieved papers that mentioned NeuroVault in the title, abstract, keywords, or body (“*neurovault*[All Fields]”) and found 194 additional collections linked to a valid publication. Altogether, we were able to identify a final sample of 1,472 NeuroVault collections that were linked to a published article with a valid DOI.

Importantly, not all collections and images in NeuroVault are “standard” collections uploaded by independent users and thus may not be suitable for meta-analysis. Specifically, 40% of the 9,502 collections were created by the Neuroscout web application (de la Vega et al., 2022), which utilizes NeuroVault to store results generated by its analysis pipelines for sharing and visualization purposes. Among the 5,756 remaining “standard” collections in NeuroVault, we examined how their images are organized based on modality, analysis level, and image type (Figure 2B). As anticipated, the majority of images in NeuroVault are categorized as “fMRI-BOLD,” with other modalities making up only 5% of the total. Interestingly, 25% of the images are not linked to existing image modalities. A surprising finding was that 88% of the images in NeuroVault are associated with individual subject analysis, while only 10% represent group-level maps. This trend is driven by a few large collections that contain many subject-level images; for instance, collections 9494, 4337, and 8996 together account for over 70,000 single-subject images. Conversely, collections that include group-level data typically contain only a handful of group-level images. Additionally, around 75% of the uploaded images are unthresholded, preserving the full richness of the data. Similarly, more than 75% of the images in NeuroVault are statistical maps (Z or T), with other types, such as variance and effect maps, representing a smaller proportion. Notably, 63% of the images are linked to valid Cognitive Atlas tasks.

### Image selection for IBMA

The primary challenge with conducting IBMA using NeuroVault is identifying relevant images for a meta-analysis. To demonstrate this process, we conducted a preliminary selection of images that could potentially be suitable for meta-analysis (Figure 3A). We focused on fMRI-BOLD images, as they are the most prevalent modality in NeuroVault, making up 70% of the total images. We specifically chose images from group-level analyses, which represent a smaller proportion compared to subject-level images, as illustrated in Figure 2B. This selection significantly reduced our sample to only 7.5% of all available images. However, the number of collections did not decrease at the same rate, reinforcing our observation that most single-subject images originate from just a few collections. Additionally, we retained only images from studies with a sample size greater than ten subjects. We selected images classified as T or Z statistics, bringing the total down to 11,422 images. Although best practices in meta-analysis suggest using meaningful units and incorporating uncertainty through standard errors, T/Z statistic maps are the most commonly shared images in NeuroVault. Moreover, we filtered for unthresholded images that covered at least 40% of the brain and were in MNI space. Finally, we narrowed our selection to 6,400 images associated with a Cognitive Atlas task, accounting for just 2.7% and 16.8% of the total image and collection sample, respectively. This selection process shows that only a small percentage of NeuroVault images are potentially relevant for image-based meta-analysis.

**Figure 3.**
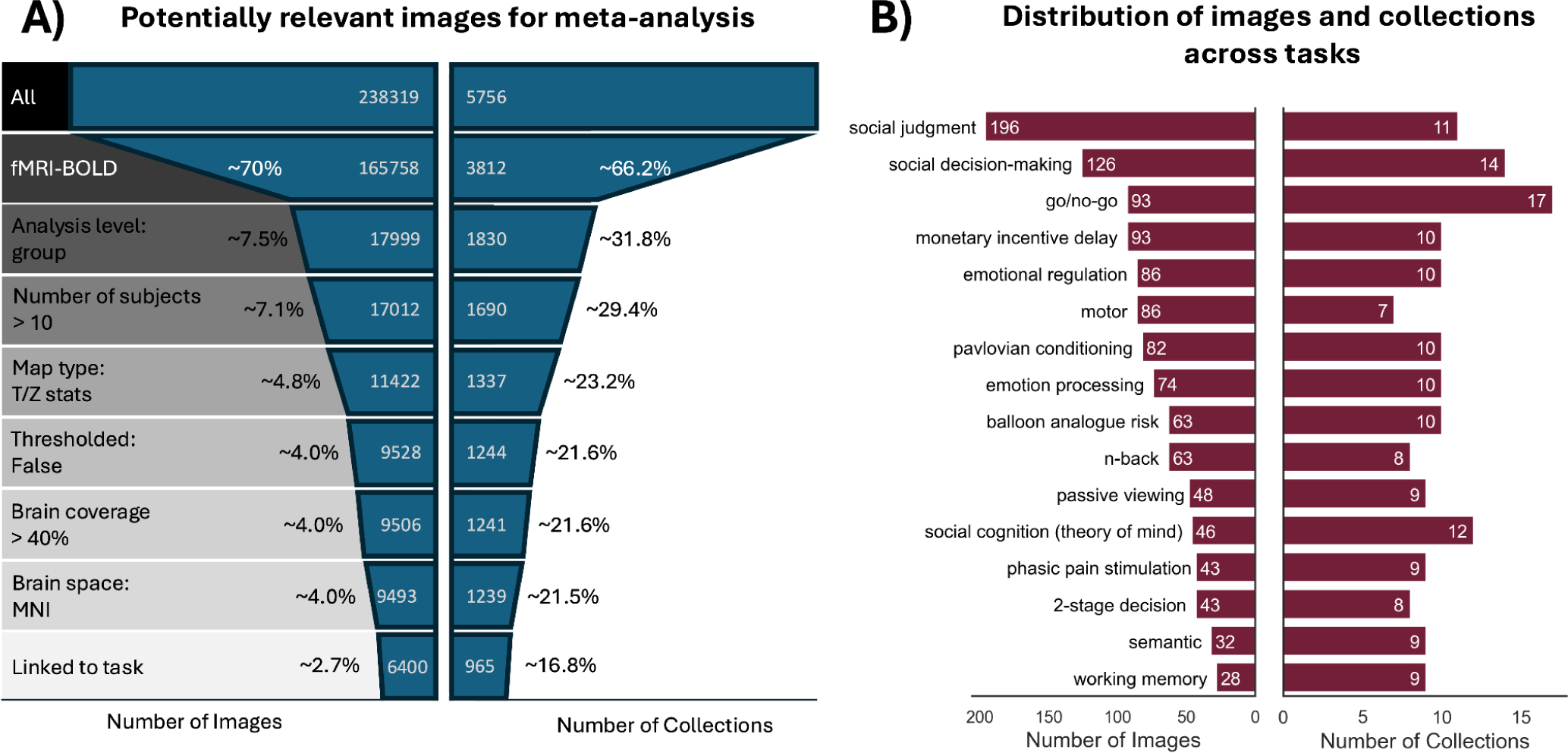
Preliminary image selection for IBMA and Cognitive Atlas task representation. **(A)** The funnel diagram presents the selection process of applying various filters to determine the sample of images suitable for meta-analysis. The number of filtered images is shown in the center on each level of the funnel, and the applied filter is presented in grayscale on the left side of the funnel. **(B)** The bar plot presents the representation of Cognitive Atlas tasks in NeuroVault, including the final set of filtered images deemed potentially usable for meta-analysis as identified by the selection in (A). The top 16 tasks are sorted by the numbers of images per task shown in the left-hand side bars and the numbers of collections per task are shown on the right-hand side.

Following that, we aimed to explore how the Cognitive Atlas tasks were represented in the remaining images and collections (Figure 3B). Social judgment and decision-making tasks had the largest number of images, with 196 and 126 images distributed across 11 and 14 collections, respectively. The go/no-go task was included in the highest number of collections, with a total of 93 images across 17 collections. Other tasks that were notably well-represented in NeuroVault included motor, emotion processing, and n-back tasks, among others. Overall, a diverse range of tasks and domains is well represented in NeuroVault, providing sufficient data for conducting image-based meta-analyses.

For a preliminary evaluation of IBMA using NeuroVault data, we selected domains whose tasks are well represented in NeuroVault. We focused on working memory, motor, and emotion processing. For working memory, we used the working memory fMRI task paradigm and n-back task. For the motor domain, we selected images linked to the motor fMRI task paradigm, motor sequencing task, and finger tapping task. Finally, for emotion processing, the emotion processing fMRI task paradigm was considered. As the reference image for the three domains, we used effect size maps from the Human Connectome Project (HCP). Additional analysis for other domains can be found in the supplementary materials. In addition to the preliminary selection shown in Figure 3A, we excluded images from the HCP NeuroVault collection. Consequently, our criteria resulted in 98 images distributed across 19 collections for working memory, 85 images in eight collections for motor, and 82 images in 14 collections for emotion processing (Table S1). This set of images is referred to as “All Images” throughout this paper.

After selecting these well-represented and exemplar domains, we applied data-driven heuristic selection methods that primarily focused on identifying images characterized by extreme values, duplicates, and inverted contrasts. For example, we only selected images with a minimum Z value greater than 1.96 and a maximum Z value less than 50. We eliminated images associated with non-statistical maps by detecting patterns in the image and file names, such as “ICA,” “PCA,” and “PPI,” among others. Consequently, we removed extreme images relative to a robust average of the whole population of images using regression slopes. This heuristic selection framework reduced the number of collections and images in domains such as working memory and emotion processing by half. However, for the motor domain, only one collection and a few images were removed (Table S1).

Finally, we manually selected the most relevant images of the publications linked in the NeuroVault collections. Our focus was primarily on the task description outlined in the paper’s method section and the specific contrast of interest. To aid in our selection, we examined the image, file name, and contrast definition fields found in the image metadata within NeuroVault. A detailed annotation of the images included/excluded during this manual selection process can be found in the supplements. Ultimately, the final sample comprised ten images from six collections for working memory, 30 images from seven collections for motor, and eight images from just two collections for emotion processing (Table S1).

### Manual image-based meta-analysis with NeuroVault

To evaluate IBMA using NeuroVault, we conducted a meta-analysis on the manually selected images corresponding to the three domains described above. The input images, initially downloaded as T/Z statistics, were converted into Cohen’s d maps using the sample sizes available in their NeuroVault metadata. We employed a baseline IBMA estimator (i.e., the mean), which performed a voxel-wise average of the input maps. For reference maps, we utilized the group-average effect size maps from the HCP S1200 data release (Barch et al., 2013; Smith et al., 2013; Uğurbil et al., 2013; Van Essen et al., 2013, 2012). Specifically, for working memory, we used the contrast “2-Back vs. 0-Back”; for motor, we aimed to reproduce the contrast representing the average of all motor movement blocks against the baseline “Motor vs. Baseline,” and for emotion processing, we employed the contrast “Face vs. Shape.” Additional details regarding these tasks and their available contrasts are described by Barch et al. (Barch et al., 2013).

We compared the results of our manual IBMA using the baseline estimator and reference maps from the HCP data. Manual IBMA effectively reproduced the reference maps for the three domains, demonstrating significant qualitative and quantitative convergence (Figure 4A). Notably, the working memory and motor domains exhibited estimated effect sizes comparable to the reference maps. In contrast, while there were similarities observed in visual brain regions for emotion processing, the average effect size estimate was weaker. Overall, these findings indicate that the NeuroVault data is suitable for IBMA.

**Figure 4.**
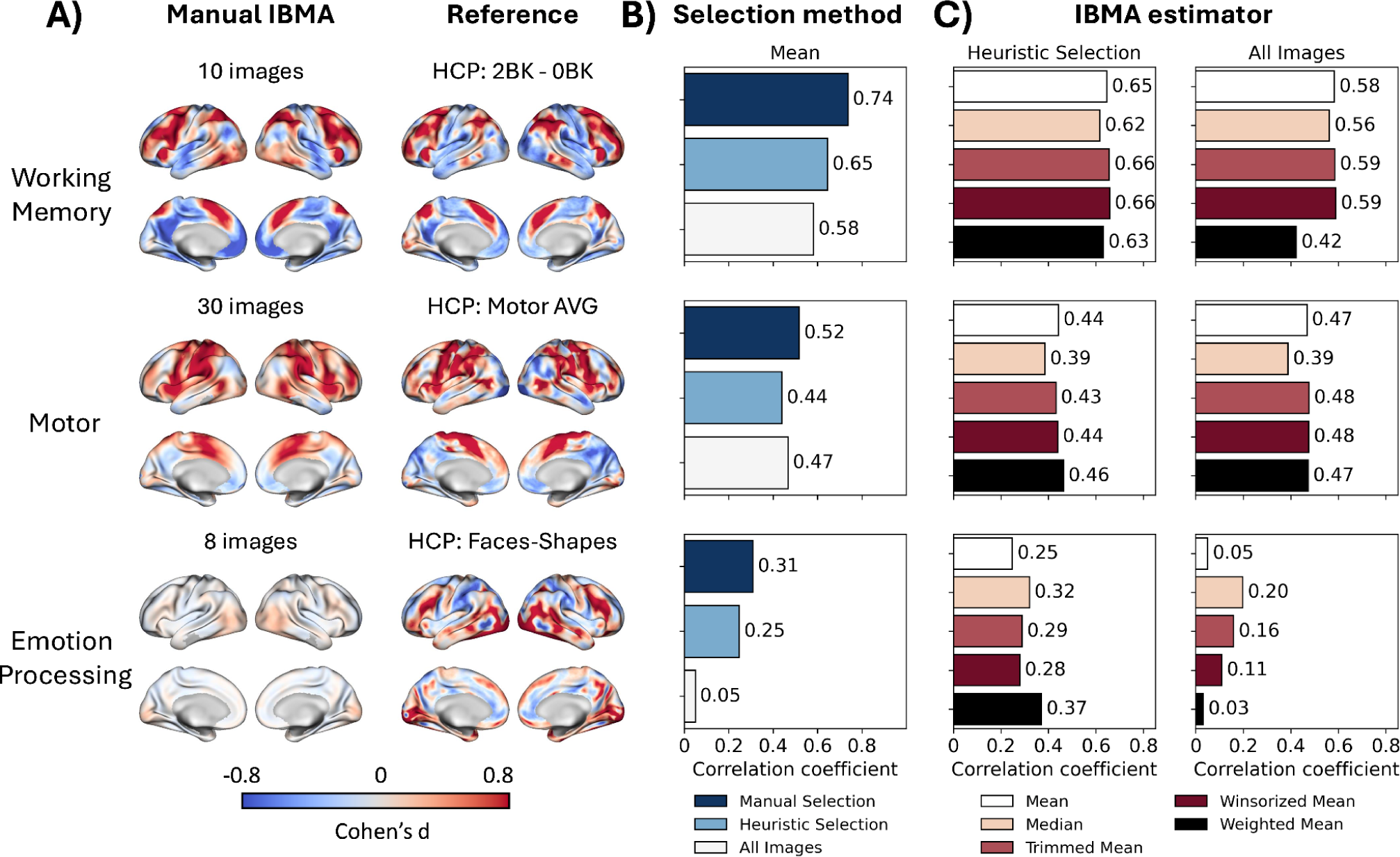
Evaluation of image-based meta-analyses with NeuroVault. **(A)** Comparison between the manual IBMAs results against the HCP’s reference maps. The top row presents the results of “working memory,” the middle row corresponds to “motor,” and the bottom row to “emotion processing.” **(B)** Comparison of the different image selection methods for the baseline estimator (i.e., mean). The bars represent the correlation with the reference maps presented in panel A. The selection methods include “Manual Selection” (navy blue), “Heuristic Selection” (light blue), and including “All Images” (light gray). **(C)** Comparison of robust estimator methods for heuristic selection of images (left-hand side plot) and including all images (right-hand side plot). The bars represent the correlation with the reference maps presented in panel A. The baseline estimator is mean (white); the robust estimators are median (apricot), trimmed mean (brick red), winsorized mean (burgundy), and weighted mean (black).

### Impact of image selection and estimator methods on image-based meta-analysis

Despite the availability of relevant data in NeuroVault, conducting a manual image-based meta-analysis still requires significant effort, as described above. Moreover, the current process for selecting relevant images from NeuroVault is not standardized, which may lead to biases introduced by researchers. With this in mind, we next considered the feasibility of applying a heuristic to selecting relevant images or including all images in a meta-analysis without the extensive manual work. In the next section, we investigate if these alternatives to the manual approach yield acceptable results.

In Figure 4B, we compared correlation coefficients from each meta-analysis based on three selection methods: (i) one that included all images from the preliminary selection, (ii) another that used a data-driven heuristic selection, and (iii) a third that relied on manual selection. As anticipated, the manual meta-analyses yielded the highest correlation with the reference maps across the three domains. Specifically, the manual IBMA for working memory showed a strong correlation of 0.74, followed by a correlation of 0.52 for motor and a weaker correlation of 0.31 for emotion processing. Surprisingly, including all images in the meta-analysis produced satisfactory results, with correlations of 0.58 for working memory and 0.47 for the motor domain. Including all images resulted in almost no correlation for emotion processing. The difference between the manual selection and the all-images approach for the motor domain was relatively small, with a 0.05 increase observed with manual selection. However, for the other two domains, the difference was more significant: there was a 0.16 gain for working memory and a 0.26 gain for emotion processing using the manual approach. Finally, the heuristic selection method showed promising results for working memory and emotion processing, suggesting that this approach effectively minimized extreme and unwanted values from the all-images sample. In fact, the correlation coefficients for these two domains were more aligned with those found through manual selection. In the case of the motor domain, the heuristic selection did not eliminate many maps and maintained the same number of collections (Table S1); as a result, the correlation achieved through the heuristic meta-analysis was slightly lower than that obtained with the all-images approach.

In addition to the baseline meta-analytic estimator, defined by the mean, we implemented four robust estimators that aimed to address extreme values in the data. For domains characterized by tasks with more homogeneous experimental paradigms, such as working memory and motor, the different estimators correlated similarly with the reference maps. Estimators that specifically target extreme values in the data, like the trimmed mean and winsorized mean, performed slightly better than the baseline estimator. However, the increase in correlation of only 0.01 does not justify the need to modify or reduce the input sample using those robust estimators. For the motor tasks, the median estimator performed worse than the other estimators, while the weighted mean showed relatively strong performance. In the case of working memory, the weighted mean yielded the best performance when including all images; this may be attributed to the presence of outlier studies with small standard errors.

For a domain defined by tasks with high heterogeneity of experimental paradigms, such as emotion processing, robust estimators significantly improved upon the baseline. Interestingly, when all images were included, we initially found a correlation of 0.05. However, using the median as a combination method increases this correlation to 0.2, which was nearly on par with the results obtained through heuristic selection using the baseline estimator. All robust estimators enhanced the correlation when applied to the heuristic selection sample, with the weighted mean achieving a correlation of 0.37, surpassing the correlation found in the baseline manual meta-analysis. Yet, the weighted mean performed the poorest when all images were considered in the meta-analysis. Overall, the trimmed mean provided the most consistent improvement across different image selection methods and domains. In manual IBMA, robust estimators performed similarly to the baseline estimator, with the mean estimator occasionally showing slightly better results. The weighted mean tended to produce the most extreme values, significantly increasing the correlation in some cases while decreasing it in others compared to the reference. Additional results of all parameter combinations tested are available in the supplementary materials.

Following this, we aimed to investigate how the estimated effect varied with different image selection methods. Figure 5 presents the distribution of estimates for the three domains, categorized by image selection methods. Not surprisingly, the results of the manual meta-analysis demonstrated broader distributions centered around zero, which indicates a relative increase in both positive and negative effect size estimates. Notably, for the motor domain, the distribution resulting from manual selection appeared more positively skewed than the other selection methods. In contrast, when we included all images in the meta-analysis, we observed the narrowest distributions among all selection methods. The distributions generated by applying the heuristic selection were slightly wider than those produced by including all images. These findings are qualitatively supported by the activation levels in the surface map plots, which clearly show that manual selection results in stronger effect estimates than the other methods. Meanwhile, the heuristic selection method still enhances the results compared to incorporating all images in the meta-analysis.

**Figure 5.**
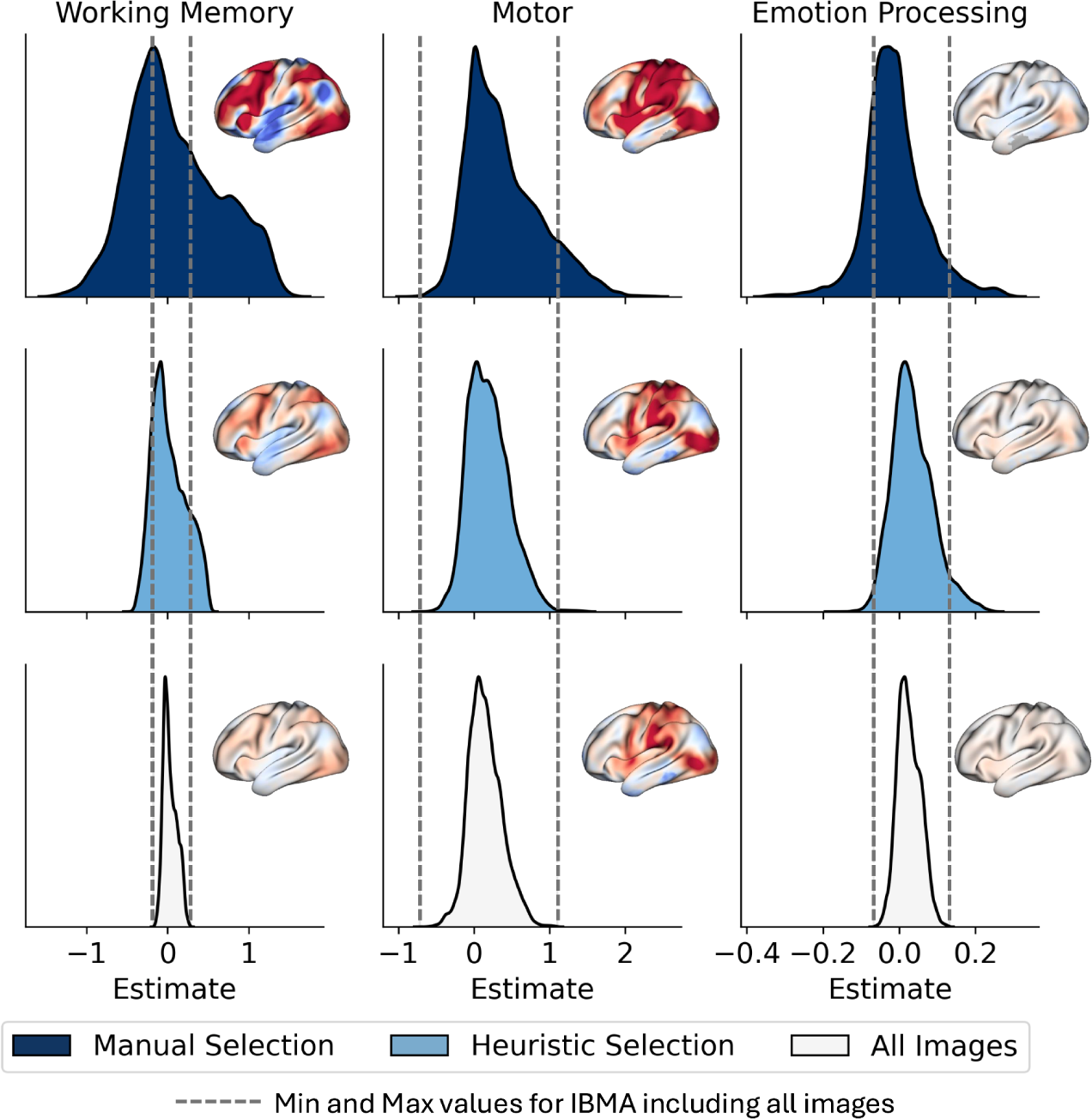
Impact of image selection on IBMA estimates. The distribution presents the value of the estimate for the baseline estimator for the three selection methods and domains. Each map’s surface brain plot color is bounded between −0.8 to 0.8. Dotted vertical gray lines delimit the minimum and maximum values in the distribution from all images as a reference to compare the distributions across selection methods. The columns represent a different domain: the leftmost column represents working memory, the middle column corresponds to motor, and the rightmost column corresponds to emotion processing. The selection methods include Manual Selection (navy blue, first row), Heuristic Selection (light blue, middle row), and All Images (light gray, bottom row).

Subsequently, we quantitatively evaluated the increased estimated values in specific areas of interest across the various domains (Figure 6). To achieve this, we focused our analysis on particular regions of interest, defined by selecting the top 10% of vertices from the reference maps (Figure 6A). The results of this evaluation for all image selection methods are displayed in Figure 6B. As anticipated, the manual meta-analysis produced the highest estimates, approaching one, for working memory and motor. Heuristic selection slightly improved the estimates compared to using all images, yielding mean estimates of 0.31 for working memory and 0.49 for motor. In contrast, including all images resulted in the lowest mean estimates across all selection methods. Interestingly, the estimates for emotion processing showed no significant differences among the selection methods, with all mean values close to zero. Regarding the meta-analytic estimators, the baseline estimator outperformed the four robust estimators for both heuristic and all image selection methods in the cases of working memory and motor. The median and weighted mean estimators exhibited the poorest performance among the five estimators. Overall, the five methods produced similar average estimates for emotion processing, with values just above zero.

**Figure 6.**
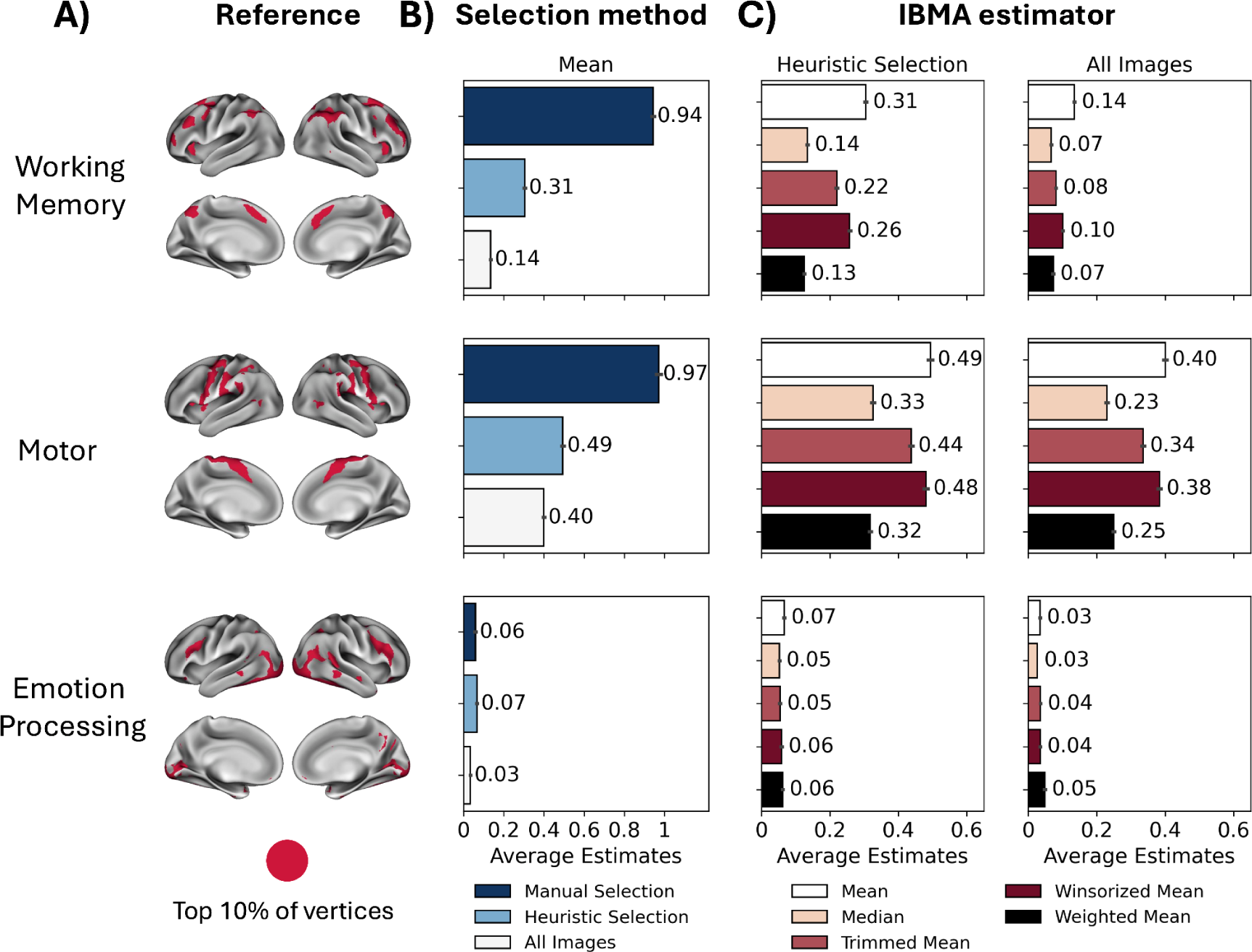
Evaluation of the size of the estimates for relevant regions of interest. **(A)** Region of interests (ROIs) highlighting the top 10 percent estimates of vertices in the cortical surface for the three domains: working memory, motor, and emotion processing. **(B)** Comparison of the different image selection methods for the baseline estimator mean. The bars represent the average estimated effect size value within the reference ROI presented in panel A. The selection methods include manual selection (navy blue), heuristic selection (light blue), and including all images (light gray). **(C)** Comparing robust estimator methods for heuristic selection of images (left-hand-side plot) and including all images (right-hand-side plot). The bars represent the average estimated effect size value within the reference ROI presented in panel A. The baseline estimator is mean (white); the robust estimators are median (apricot), trimmed mean (brick red), winsorized mean (burgundy), and weighted mean (black).

## Discussion

The NeuroVault repository currently houses a vast collection of brain images. However, the absence of standardized methods, accessible tools, and clear guidelines has hindered the reusability of this rich data source for secondary data analyses, such as IBMA. In this study, we examined the current state of the NeuroVault data, developed an automatic heuristic for selecting images, implemented five estimation methods, and provided guidelines for conducting IBMA. First, we highlighted the rapid growth of NeuroVault as a comprehensive repository for sharing neuroimaging statistical images. We noted a diverse representation of tasks and domains supporting secondary data analyses. Second, we demonstrated the feasibility of conducting IBMA with NeuroVault by reproducing reference maps from the HCP data for three distinct domains: working memory, motor, and emotion processing. We emphasized the importance of proper image selection for IBMA, particularly for heterogeneous domains such as emotion processing. Finally, we implemented robust IBMA estimators to manage extreme or spurious images within the input sample, underscoring their relevance for more complex domains.

NeuroVault has been widely adopted by researchers in the neuroimaging community. In just ten years, it has grown from 61 images at its launch in 2013 to over a quarter of a million images today. Our systematic analysis revealed significant challenges within the NeuroVault repository. Despite hosting over 238,000 images across 5,756 collections, only a small fraction were potentially relevant for image-based meta-analysis (IBMA). Specifically, only 2.7% of the images met our preliminary inclusion criteria, where requiring group-level images for meta-analysis represented the strictest criteria. This was expected since the subject-level data constitutes 88% of the total images. However, we noticed that most of those individual subject data were concentrated in a few large collections. As a result, we ultimately considered a reasonable representation of NeuroVault collections. The plateau shown in Figure 2B in publication-linked collections in recent years may suggest a potential decline in the availability of imaging data for meta-analysis in the future. However, we believe this pattern is likely due to publication time lags, and we expect that more existing collections will eventually be linked to publications. While some collection owners may not add the publication DOI to their collections post-publication, we can effectively search PubMed for any publications missing from the NeuroVault collections. In summary, our results address previous concerns about the feasibility of IBMA concerning data availability. We have shown that NeuroVault contains enough data to conduct IBMA despite strict filtering criteria. The images suitable for meta-analysis cover a diverse range of domains, including social judgment, decision-making, motor, emotion processing, and working memory.

To demonstrate the feasibility of using IBMA with NeuroVault data, we analyzed three domains that have been extensively studied using fMRI and for which there are reference maps available in the HCP dataset. These domains included working memory, motor, and emotion processing. Our manual meta-analysis produced maps that closely resembled the reference maps for each of these domains. We highlighted the significant potential of NeuroVault for future large-scale image-based meta-analyses, particularly as the repository expands and data-sharing practices improve through standardized uploading procedures.

Our comprehensive analysis of the NeuroVault repository revealed that many collections and images face data quality issues, such as incorrect annotations, missing publications, and non-statistical maps. This underscores the urgent need for standardized data curation methods to facilitate secondary analysis using NeuroVault data. While we initially conducted a manual meta-analysis to address data quality concerns, this approach is labor-intensive. Additionally, the process of manually selecting relevant images lacks standardization, which can lead to inconsistencies and reproducibility issues across studies. To tackle these challenges, we developed and validated an automated framework for removing potentially spurious images and other outliers. This framework allows researchers to efficiently identify and filter suitable statistical maps for IBMA, thereby promoting good practices and guiding future meta-analyses. Our method has enhanced reliability and reproducibility, and case studies demonstrated how effective data filtering and model selection can significantly influence meta-analytic results.

Our findings indicate that, although manual image selection is considered the gold standard for image-based meta-analysis (IBMA), alternative approaches show promising potential for enhancing this process using NeuroVault. Manual selection consistently produced the strongest correlations with reference maps across all domains and yielded the highest effect size estimates. The performance gap between manual and automated approaches varied depending on the domain. In domains with more homogeneous tasks, such as motor, the results of including all images were comparable to those obtained through manual selection. However, manual selection resulted in larger positive estimated effect sizes. For more complex domains, like emotion processing, manual selection demonstrated significant advantages, showing a 0.26 improvement in correlation compared to using all images. Interestingly, the estimated effect sizes did not show significant differences across the selection methods in this case. Surprisingly, our heuristic selection method achieved correlations closer to those obtained through manual selection for working memory and emotion processing while also showing modest improvements in effect size estimates compared to using all images. These results suggest that while manual selection remains optimal for detecting stronger effect sizes and achieving broader activation distributions, combining heuristic selection with appropriate estimators could provide a practical balance between accuracy and efficiency in image-based meta-analysis.

The effectiveness of robust estimators varied by domain, with the trimmed mean displaying consistent performance across different selection methods. In particular, for emotion processing, robust estimators significantly improved correlations when using all images, with the median estimator increasing correlation from 0.05 to 0.20. However, for working memory and motor, the baseline estimator outperformed robust estimators when using heuristic or all-image selection methods. Importantly, no one “best” method for aggregating images in a meta-analysis exists. The median is the most robust method but the most variable. Winsorization comes closest in efficiency to the mean but regards outlying observations as conveying useful information; trimming is less efficient than winsorization but completely removes the influence of extreme observations. In general, we are most confident in our results when all these robust methods give similar answers and agree with the conventional mean (e.g., the results for the heuristic selection for working memory and motor). When these robust methods dramatically differ from the mean (e.g., the results of including all images for emotion processing), this indicates that extreme observations play an appreciable role. Unfortunately, though, the reduced efficiency implies that with a very small sample size, these robust methods could produce rather erratic results.

Several limitations in the current work must be highlighted to move the field forward. The meta-analytic estimator method used in this work solely focuses on handling extreme and unwanted observations. First, we considered all images to be independent of each other. Nevertheless, in most cases, NeuroVault images belong to the same paper or collection. Thus, they cannot be considered independent since they may undergo the same analytic pipeline or could correspond to the same population. Critically, our assumption of independence can lead to inflated false positive results. A recent work on same data meta-analyses has studied this issue, proposing multiple models to account for dependencies (Lefort-Besnard et al., 2024). These models were evaluated for Stouffers and the generalized least square problem. Future work is required to incorporate and test these models in our robust estimator methods. Second, we only included analysis for three domains as a proof of concept. Although additional analyses were conducted and presented as supplementary information, such results could not be thoroughly evaluated, given the lack of reference maps from large-scale fMRI studies. Moreover, other domains could not be considered because of data availability. However, we are confident that, as NeuroVault repositories continue to grow, our observations will be evaluated at a larger scale. The results presented in this paper, along with the methodology and tools, will be critical in extracting meaningful scientific insights from this increasingly large and complex database.

Taken together, this work is the first to present a systematic approach for implementing IBMA using the NeuroVault repository. Our findings indicate that NeuroVault currently has sufficient data to conduct IBMA for a diverse range of tasks and domains. Although we encountered several data quality issues, we successfully reproduced reference maps from the HCP for three distinct domains using a simple heuristic selection method. For domains involving standardized experimental task paradigms, such as motor and working memory, utilizing the baseline mean estimator along with all available images or a heuristic selection yielded reasonable results. However, for domains characterized by heterogeneous task paradigms, such as emotion processing, it is essential to employ a robust estimator and undertake a heuristic or manual selection of images. Most importantly, we have provided methods and guidelines to support future meta-analyses utilizing NeuroVault data.

## Methods

### Databases NeuroVault

NeuroVault (https://neurovault.org) is a web-based repository of fMRI statistical maps from neuroimaging studies (Gorgolewski et al., 2015). The brain maps are grouped in collections that are created and updated voluntarily. This repository can be explored and downloaded with the help of an API, which is supported by some Python neuroimaging tools (e.g., Nilearn and NiMARE). As of January 2024, NeuroVault contained 238,319 maps distributed in 5,756 collections, of which approximately 1,473 were associated with a published paper. The collections were linked to papers using the DOI field and collection description from their metadata. We also searched PubMed for articles that matched the collection’s title. Additionally, we conducted an extensive search using Pubget, an open-source Python tool for collecting data for biomedical text mining (https://neuroquery.github.io/pubget/pubget.html). We performed a query and retrieved papers that mentioned NeuroVault in the title, abstract, keywords, and body of the articles (“*neurovault*[All Fields]”).

To explore the NeuroVault database, we created an SQL query and exported the database contents to human-readable tables while filtering sensitive user information. This provided sufficient metadata from all collections and images to investigate the entire database without downloading the files. The images identified as usable for IBMA (see the following section on the image selection framework) were downloaded along with their metadata and converted to a NiMARE Dataset object to leverage existing IBMA methods implemented in NiMARE.

### Cognitive Atlas

Cognitive Atlas (Poldrack et al., 2011) (https://www.cognitiveatlas.org/) is an online repository of cumulative knowledge from experienced researchers from the psychology, cognitive science, and neuroscience fields. The repository currently offers two knowledge bases: 907 cognitive concepts and 841 tasks with definitions and properties. Cognitive concepts contain relationships with other concepts and tasks, with the goal of establishing a map between mental processes and brain function. It provides an API to download the database, which is also integrated into NiMARE.

### Image selection framework

#### Preliminary selection

Using the available metadata from the retrieved tables, we set different inclusion criteria for images to be considered for a meta-analysis. We focused on fMRI-BOLD images, as they are the most prevalent modality in NeuroVault. Note that the methods presented in this paper should work with other image modalities (e.g., PET, diffusion MRI, structural MRI). Still, only fMRI-BOLD had enough data in NeuroVault for meta-analyses. Then, we specifically chose images from group-level analyses. Additionally, we retained only images from studies with a sample size greater than ten subjects. Next, we selected images classified as T or Z statistics. Although best practices in meta-analysis suggest using meaningful units and incorporating uncertainty through standard errors, T/Z statistic maps are the most commonly shared images in NeuroVault (Maumet and Nichols, 2016). We discuss this further in the following sections. Upon review, it is important to note here that plenty of images in NeuroVault are labeled as “Other” for the image type. Nonetheless, most of those images actually correspond to known image types (e.g., T/Z statistic). As a result, we relabeled those images to their original type if keywords such as “zstat,” “tstat,” “Z_,” or “T_” were present in the image name, file name, or image description. Following that, we retained unthresholded images that cover 40% of the brain and are in MNI space. Ultimately, we narrowed our selection to images associated with a Cognitive Atlas task.

#### Heuristic selection

Even after applying the previous strict preliminary inclusion criteria, we still found plenty of wrongly annotated images, especially representing other image modalities and others with extreme values. Therefore, we developed an automatic heuristic selection to remove those spurious images from the meta-analysis. The heuristic selection consisted of two steps. First, we removed all images from collections that lacked a link to a publication. Also, images with a minimum Z value smaller than 1.96 (i.e., Z score for a 0.05 p-value) were removed as they potentially consisted of mislabeled correlation maps, inverted p-value maps, or did not contain voxels statistically significant. We also excluded images with a maximum Z score larger than 50. Although the number 50 is arbitrary, we wanted to detect images with an unusually large signal. For example, mislabeled BOLD or COPE (contrast of parameter estimates) images or others resulting from studies with a huge sample size. Additionally, using the image metadata, we analyzed the image and file name. We removed those containing keywords such as “ICA,” “PCA,” “PPI,” “seed,” “functional connectivity,” “cope,” “tfce,” and “correlation,” which represent modalities not of interest for the meta-analysis of the current work.

Second, we wanted to detect and remove extreme images relative to a robust average of the whole population of images. Note that the population of images to calculate the average was considered on a domains basis, and not the entire sample of images from NeuroVault. After creating the robust average images (i.e., the median), we made a rough segmentation of ‘signal’ and ‘noise’ voxels. For signal, we defined the mask as the bottom and top 10% of voxels (by rank order); for ‘noise,’ conversely, we selected the 20% of voxels with the smallest magnitude (i.e., closest to 0). Then, we performed the correlation exercise only on the ‘signal’ voxels (i.e., the correlation *r_iM_* between each image *i* and the median *M*, only in signal voxels). We calculated the standard deviation among noise voxels for each image *S*_*i*_ and the median *S*_*M*_. Ultimately, the regression slope *Slope_i_* for each image (*i*) relative to the median image was determined by:

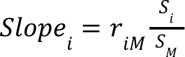

It is quite common for NeuroVault users to upload inverted contrasts and duplicates. For example, one might find two images representing the same contrast (such as House > Face) but with the signs reversed (i.e., Face > House). This creates problems for meta-analyses, as these images effectively cancel each other out when aggregated. Additionally, it is typical for users to upload multiple images of the same contrast, differing only by the covariate used in the group-level analysis. These can be considered duplicates, especially when the covariate does not influence the final estimate. To identify duplicates, we utilize the correlation matrix of the input samples. Image pairs with a correlation close to 1 are considered duplicates, while those with a correlation close to −1 are labeled as inverted contrasts. From the identified duplicates, we randomly selected one image from each pair. For pairs of inverted contrasts, we choose the image with a positive slope relative to the median image.

Finally, we removed images with extreme regression slope values relative to the population median using the interquartile range (IQR) method. First, we sorted the slopes in ascending order and calculated the first and third quartiles (Q1 and Q3). The IQR was defined as the difference between Q3 and Q1 (IQR = Q3 - Q1). Next, we determined the lower and upper bounds: the lower bound is calculated as Q1 - 1.5 * IQR, and the upper bound as Q3 + 1.5 * IQR. We compared each slope to these bounds, removing any images with slopes that were smaller than the lower bound or larger than the upper bound.

#### Manual selection

The manual meta-analysis served as an initial evaluation of the IBMA with NeuroVault data, as manually selected samples are less likely to include spurious or non-relevant images. Our primary focus was on the task description outlined in the method section of the paper, as well as the specific contrast of interest. To assist in our selection, we examined the image metadata in NeuroVault, specifically the image, file name, and contrast definition fields. For instance, in the case of a domain involving working memory tasks like the n-back task, we reviewed the paper associated with the collections containing images from this task. Images related to the cognitive atlas task were identified using the metadata field “cognitive_paradigm_cogatlas.” We prioritized the section of the paper that describes the task used in the study and checked whether the contrast of interest for the meta-analysis (e.g., 2-back vs. baseline) was present. If the study did explore this contrast, we then examined all images available in the corresponding NeuroVault collection that met our preliminary and heuristic selection criteria. To locate the relevant images, we searched for the contrast of interest in various fields of the image metadata, including the Cognitive Atlas contrast (“cognitive_contrast_cogatlas”), image title, file name (under the “file” field in NeuroVault), and contrast definition (found in the “contrast_definition” field in NeuroVault).

### Image-based meta-analysis

A preliminary evaluation of IBMA using NeuroVault data, was performed on domains whose tasks are well represented in NeuroVault. We focused on working memory, motor, and emotion processing. For working memory, we used the working memory fMRI task paradigm, and n-back task. For the motor domain, we selected images linked to the motor fMRI task paradigm, motor sequencing task, and finger tapping task. Finally, for emotion processing, emotion processing fMRI task paradigm. As the reference image for the three domains, we used effect size maps from the Human Connectome Project (HCP) Stouffer’s method is the most popular approach for combining individual images in IBMA (Stouffer et al., 1949). This method combines test statistics, assuming that the input values are standardized to have a mean of zero and a variance of one (i.e., Z scores) (Camille and Thomas, 2014). However, best practices in meta-analysis suggest using values with meaningful units, incorporating uncertainty through standard errors instead of relying solely on Z or T statistics (Salimi-Khorshidi et al., 2009). A significant challenge in neuroimaging, particularly with functional fMRI derivatives data, is that researchers typically share only T or Z statistic maps instead of actual estimates with their associated standard errors. Moreover, even if estimates were provided, we often lack information about the units of measurement for these estimates. For instance, the FSL fMRI pipeline scales mean brain intensity to 10,000 (Jenkinson et al., 2012), while the SPM pipeline targets a scale of 100 (Friston et al., 2007), which usually aligns more closely with a value around 130. To address these concerns, we used standardized effect size as input for meta-analyses, recently proposed by Bossier, Nichols, and Moerkerke (2019) (Bossier et al., 2019). The use of standardized effect sizes, which have no units, it facilitates aggregating results from different studies.

We reconstructed the standardized effect size (i.e., Cohen’s d) from the Z/T score maps using the sample size available in the image metadata in NeuroVault and assuming that a one-sample t-test was used. It is important to note that various types of analyses exist, such as two-sample analyses that compare different groups, correlations that assess the relationship between brain response and a covariate, and F-tests that compare two or more groups, among others. However, since most task-based fMRI studies utilize a one-sample t-test, this assumption is reasonable unless we have more specific information about each study.

By definition, the population standardized effect size for a one-sample analysis is the population mean divided by the population standard deviation (*d* = µ/σ). For *N* subjects, we can thus compute an estimate of standardized effect sizes from a one-sample t-test t as

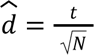

While the sample mean is unbiased for the population mean, the sample Cohen’s d above is not unbiased for the population *d*. A bias correction due to Hedges is

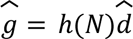

where

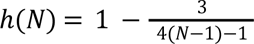

In the following sections, we consider the effect estimate *y_kv_* as the Hedges’ g estimate:

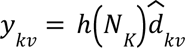

Let *y_kv_* be the effect estimate for contrast *k* at voxel *v*, *k* = 1, …, *K*, *v* = 1, …, *V*; denote the corresponding standard error be *s*_*kv*_, and let *N*_*k*_ be the sample size.

### Baseline estimator

#### Mean

The baseline estimator used in this work is the mean, where input images are aggregated using the equation for each voxel:

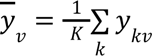

#### Robust meta-analysis methods

Meta-analysis methods can be significantly affected by extreme observations. In most cases, except for random-effects meta-analysis, an extreme data point is treated as strong evidence of an effect and can skew the results towards significance. In a random-effects meta-analysis, however, an outlier can increase heterogeneity and ultimately reduce the significance of the findings. In this context, we proposed four robust alternative methods to the baseline mean. These methods are designed to tolerate a certain fraction of corrupted data while providing reasonable estimates of the overall effect. Although we will only define these methods for unit-based effects (such as Cohen’s d or Hedge’s g), they can also be applied analogously to test statistics.

#### Median

The first robust method, the median, is denoted for a given voxel v as 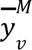. The median has a “breakdown point” of 50%, meaning it can remain unbiased even if up to 50% of the data is corrupted. Unlike the sample mean, the standard error of the median is affected by the actual distribution of the data. Therefore, the following result, unfortunately, relies on the assumption of normality:

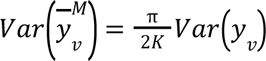

While concerns about outliers are valid, it is also essential to consider the variance of the data (denoted as *Var y_v_*). In this regard, It is essential to recognize that the mean is generally more efficient than the median. When considering variance, the mean of K values has a variance that is 1/K times that of the original data. In contrast, the variance of the median is π/2, approximately 1.57 times larger. As a result, the standard deviation for the median is roughly 25% greater than the mean’s.

#### Trimmed and Winsorized mean

The two other robust methods considered in this work are the Trimmed mean and the Winosorized mean. For these methods, we need additional notation: Let *y*_(1)*v*_ ≤ ⋯ ≤ *y*_(*k*)*v*_ ≤ ⋯ ≤ *y*_(*K*)*v*_ be the ordered

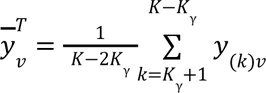

For Winsorization, the extreme values are replaced with their nearest neighbors.

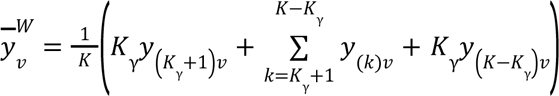

Regarding the choice of *γ*, we want to consider removing at least two studies, one from each tail, and so *γ* = 2/*K*. When there are many studies, there is no principled here one heuristic: If *K* = 20, then the minimum is *γ* = 2/20 = 10%, suggesting *γ* = 10% in general. Fortunately, for normally distributed data, the tails are so light that trimmed and winsorized mean with *γ* = 10% or even *γ* = 20% have good efficiency, though, with small *K*, it is probably best to stick to the lower value of 10%.

#### Weighted Mean: Fixed effect Hedges’ g

The weighted mean through a fixed effect meta-analysis is another robust estimator, where studies with smaller standard errors are weighted higher in the average of the input data. Using the equation for a fixed effect meta-analysis, assuming 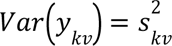, with the optimal weights 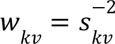, the fixed effect meta-analytic estimate is

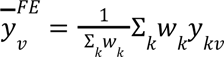

Where Bossier, Nichols, and Moerkerke (2019) defined the standard error of bias-corrected Cohen’s d, i.e. Hedge’s g, as

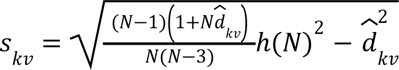

#### Evaluating image-based meta-analyses

We assess the meta-analyses with all different combinations of parameters (i.e., image selection method and estimator approach) with the help of reference images from the task-fMRI group-average effect size maps from the HCP S1200 data release (Barch et al., 2013; Smith et al., 2013; Uğurbil et al., 2013; Van Essen et al., 2013, 2012). Specifically, we used the contrast of 2-back versus 0-back for working memory. For the motor domain, we used the contrast representing the average of all motor movement blocks against the baseline in the HCP task. The contrast “Face vs. Shape” was our reference for emotion processing. Additional details regarding these tasks and their available contrasts can be found here (Barch et al., 2013).

The comparisons between our IBMA results and the reference maps focused on evaluating image similarity and increased estimates in specific brain regions of interest. To quantitatively evaluate the similarity of the images, we calculated correlation coefficients between vertex-level unthresholded meta-analytic estimate maps from the IBMA and the reference unthresholded group-average effect size maps from the HCP. Since the reference maps were defined in CIFTI format, containing all grayordinates from the subcortical structure and cortical regions, we transformed the meta-analytic maps from the MNI152 space to the standard MNI fsLR 32K 2-mm mesh surface space of the HCP, using the `mni152_to_fslr` function from the Neuromaps’ transforms module (Markello et al., 2022). For simplicity, we focused only on cortical regions for our evaluation. Subsequently, we quantitatively evaluated the increased estimated values in specific areas of interest across the domains. To achieve this, we focused our analysis on particular regions of interest, defined by selecting the top 10% of vertices for each reference map.

## Conclusions

This study advances neuroimaging research by providing a comprehensive, reproducible framework for conducting IBMA using NeuroVault data. Our findings highlight both the challenges and potential of the NeuroVault repository. The methodology presented here offers researchers a robust set of tools and methods for assessing data quality, implementing flexible image selection strategies, and conducting reliable meta-analyses across diverse domains. Future work should focus on further refining automated selection techniques, expanding the range of domains analyzed, and developing more sophisticated robust estimator methods. As the NeuroVault repository continues to grow, such standardized approaches will be critical in extracting meaningful scientific insights from the increasingly large and complex neuroscience literature.

## Acknowledgments

Special thanks to the FIU Instructional & Research Computing Center (IRCC, http://ircc.fiu.edu) for providing the HPC and computing resources that contributed to the research results reported in this paper.

## Ethical Statement

The Human Connectome Project provided the ethics and consent needed for the study and dissemination of HCP data. This secondary data analysis was approved by the Institutional Review Board of Florida International University.

## Data Accessibility

The functional connectivity data were provided by the Human Connectome Project, WU-Minn Consortium (Principal Investigators: David Van Essen and Kamil Ugurbil; U54-MH091657) funded by the 16 NIH Institutes and Centers that support the NIH Blueprint for Neuroscience Research; and by the McDonnell Center for Systems Neuroscience at Washington University.

The imaging data for meta-analyses used in this project are publicly available for download at https://neurovault.org/.

## Code Availability

This project relied on multiple open-source Python packages, including: Jupyter (Kluyver et al., 2016), Matplotlib (Hunter, 2007), Neuromaps (Markello et al., 2022), NiBabel (Brett et al., 2020), Nilearn (Abraham et al., 2014), NiMARE (Salo et al., 2024, 2023), PyMARE (Yarkoni et al., 2024), NumPy (van der Walt et al., 2011), Pandas (McKinney, 2010), PtitPrince (github.com/pog87/PtitPrince), PySurfer (Waskom et al., 2020), Scikit-learn (Pedregosa et al., 2011), SciPy (Virtanen et al., 2020), Seaborn (Waskom, 2021), and SurfPlot (Gale et al., 2021),. We also used the HCP software Connectome Workbench (wb_command version 1.5.0, (Marcus et al., 2011)).

All code required to reproduce the analyses and figures in this paper is available on GitHub at https://github.com/NBCLab/large-scale-ibma. All data and resources that resulted from this paper (e.g., connectivity gradients and trained meta-analytic decoders) are openly disseminated and made available on the Open Science Framework (OSF) at https://osf.io/w7zcp/, including the links to the GitHub repository, and figures.

## Author Contributions

ARL, JAP, AdlV, JBP, TEN, and JDK conceived and designed the project. JAP, AdlV, RWB, and JDK analyzed data. JAP, RWB, and JDK contributed scripts and pipelines. JAP, TEN, and ARL wrote the paper, and all authors contributed to the revisions and approved the final version.

