## Supplemental Information for "Advancing image-based meta-analysis for fMRI: A framework for leveraging NeuroVault data"

**Table S1: Distribution of collections and images across selection methods by domains.**

| Domain | Image selection method | Number of collections | Number of images |
|---|---|---|---|
| Working Memory | All images | 19 | 98 |
| | Heuristic | 11 | 51 |
| | Manual | 6 | 10 |
| Motor | All images | 8 | 85 |
| | Heuristic | 8 | 73 |
| | Manual | 7 | 30 |
| Emotion Processing | All images | 14 | 82 |
| | Heuristic | 7 | 39 |
| | Manual | 2 | 8 |
| Pain | All images | 14 | 88 |
| | Heuristic | 11 | 41 |
| | Manual | 7 | 13 |
| Social Cognition | All images | 33 | 153 |
| | Heuristic | 14 | 47 |
| | Manual | 8 | 11 |
| Response Inhibition | All images | 20 | 99 |
| | Heuristic | 15 | 61 |
| | Manual* | - | - |
| Reward & Decision Making | All images | 15 | 109 |
| | Heuristic | 9 | 61 |
| | Manual* | - | - |
| Risk | All images | 15 | 70 |
| | Heuristic | 9 | 43 |
| | Manual* | - | - |
| Visual Perception | All images | 9 | 48 |
| | Heuristic | 8 | 40 |
| | Manual* | - | - |

*Manual meta-analysis was not performed.

### Image-based meta-analyses

#### Working memory

- Working memory fMRI task paradigm
- N-back task

Contrast of interest: 2-back vs baseline; 2-back vs 0-back

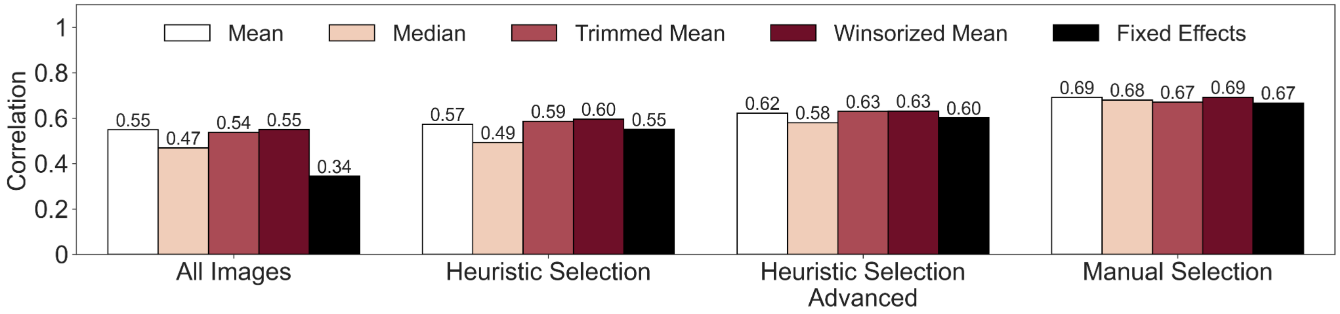


**Figure S1. Evaluation of working memory image-based meta-analyses with NeuroVault.** Comparison of the different image selection methods and estimators. The bars represent the correlation with the reference maps.

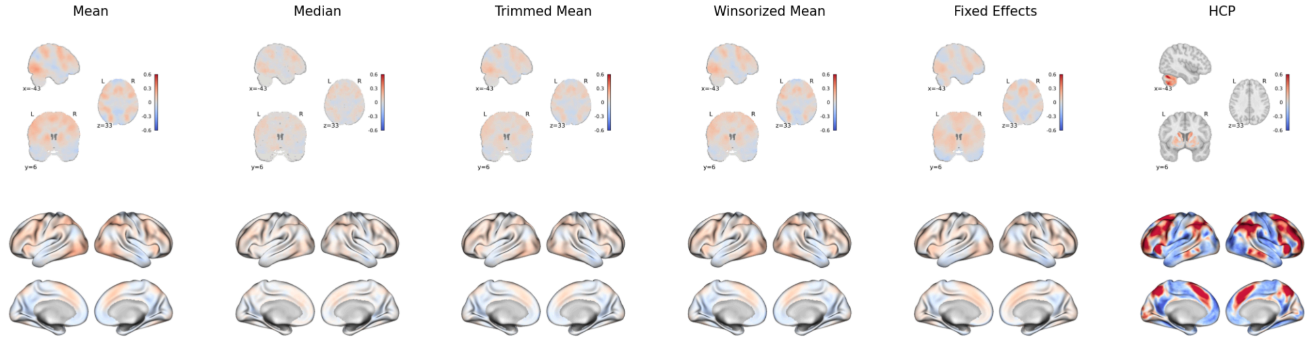


**Figure S2. IBMA result for working memory, including all images.**

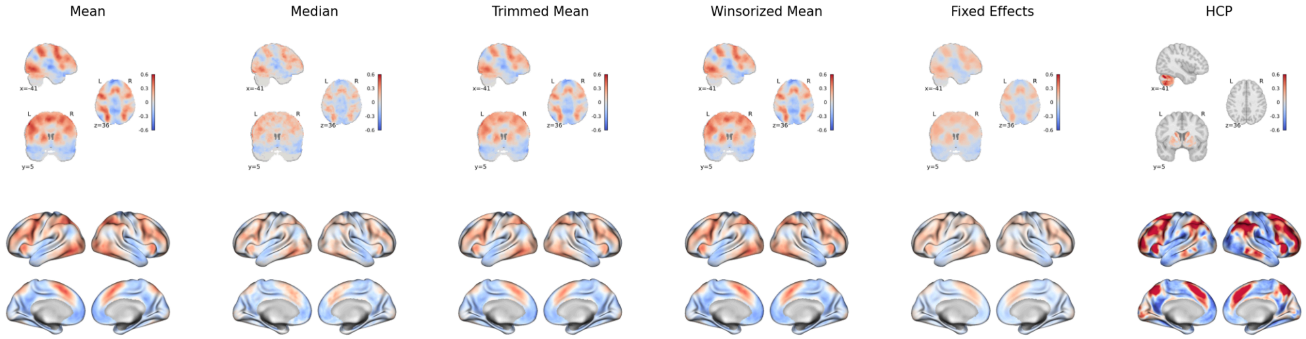


**Figure S3. IBMA result for working memory, with automatically selected images.**

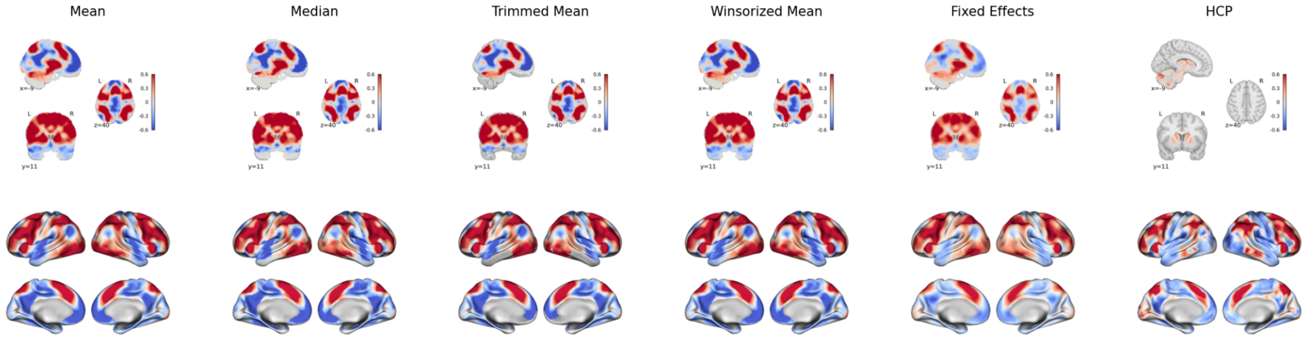


**Figure S4. IBMA result for working memory, with manually selected images.**

#### NeuroVault Collections and PMIDs

### 2230

- Link: https://neurovault.org/collections/2230
- Paper: https://pubmed.ncbi.nlm.nih.gov/28288700
- Include: True
- Images:
  - https://neurovault.org/images/42803
    - Include
  - https://neurovault.org/images/42805
    - Include
  - https://neurovault.org/images/42807
    - Include
  - https://neurovault.org/images/42810

  - https://neurovault.org/images/42811
  - https://neurovault.org/images/42812

**3191**

- Link: https://neurovault.org/collections/3191
- Paper: https://pubmed.ncbi.nlm.nih.gov/29521161
- Include: False
  - This paper corresponds to theory of mind (ToM) instead of working memory. ONly include 1-back contrasts
- Images:
  - https://neurovault.org/images/57444
  - https://neurovault.org/images/57446
  - https://neurovault.org/images/57447
  - https://neurovault.org/images/57464

**3373**

- Link: https://neurovault.org/collections/3373
- Paper: https://pubmed.ncbi.nlm.nih.gov/29860081
- Include: False
  - This paper has a task different that a classic n-back task: a human WM study in the tactile modality where participants had to memorize the spatial layout of patterned Braille-like stimuli presented to the index finger.
- Images:
  - https://neurovault.org/images/59429

**3192**

- Link: https://neurovault.org/collections/3192
- Paper: https://pubmed.ncbi.nlm.nih.gov/30150393
- Include: False
  - The images reported in the paper correspond to PCA of the fMRI data
- Images:
  - https://neurovault.org/images/57498
  - https://neurovault.org/images/57499

**3264**

- Link: https://neurovault.org/collections/3264
- Paper: https://pubmed.ncbi.nlm.nih.gov/30221091
- Include: True

   - Not an n-back task, but one contrast is WM
- Images:
   - https://neurovault.org/images/58189
   - https://neurovault.org/images/58190
   - https://neurovault.org/images/58191
   - https://neurovault.org/images/58192
      - Include

**4623**

- Link: https://neurovault.org/collections/4623
- Paper: https://pubmed.ncbi.nlm.nih.gov/30608970
- Include: True
   - Contrast in paper 2-back vs baseline
- Images:
   - https://neurovault.org/images/109900
   - https://neurovault.org/images/109901
      - Selected too: 2-back > 1-back
   - https://neurovault.org/images/109902
      - Image selected for meta-analysis: 2-back vs baseline (identify in the image name)

**4742**

- Link: https://neurovault.org/collections/4742
- Paper: https://pubmed.ncbi.nlm.nih.gov/30905834
- Include: True
   - Not an n-back task, but selective working memory was tested with attended > ignore
- Images:
   - https://neurovault.org/images/111340
   - https://neurovault.org/images/111344

**4998**

- Link: https://neurovault.org/collections/4998
- Paper: https://pubmed.ncbi.nlm.nih.gov/31954729
- Include: False
   - Not an n-back task
- Images:
   - https://neurovault.org/images/123496
   - https://neurovault.org/images/123497
   - https://neurovault.org/images/123498
   - https://neurovault.org/images/123499

- https://neurovault.org/images/123500
- https://neurovault.org/images/123501
- https://neurovault.org/images/123502
- https://neurovault.org/images/123503
- https://neurovault.org/images/123504
- https://neurovault.org/images/123505
- https://neurovault.org/images/123506
- https://neurovault.org/images/123507
- https://neurovault.org/images/123508
- https://neurovault.org/images/123509

**8710**

- Link: https://neurovault.org/collections/8710
- Paper: https://pubmed.ncbi.nlm.nih.gov/34089873
- Include: False
    - Not an n-back task
- Images:
    - https://neurovault.org/images/405163

**9469**

- Link: https://neurovault.org/collections/9469
- Paper: https://pubmed.ncbi.nlm.nih.gov/34738280
- Include: True
    - Contrast of interest 2-back vs 0-back
- Images:
    - https://neurovault.org/images/442124
    - https://neurovault.org/images/442125
    - https://neurovault.org/images/442126
        - Image selected for meta-analysis: 2-back vs 0-back (identify in the image name)
    - https://neurovault.org/images/442127
    - https://neurovault.org/images/442128
    - https://neurovault.org/images/442129
    - https://neurovault.org/images/442130
    - https://neurovault.org/images/442131
    - https://neurovault.org/images/442132
    - https://neurovault.org/images/442133

**13042**

- Link: https://neurovault.org/collections/13042
- Paper: https://pubmed.ncbi.nlm.nih.gov/35511160
- Include: True
- Images:
    - https://neurovault.org/images/787480
        - Image selected for meta-analysis: 2-back vs 0-back (identify in the contrast definition)
    - https://neurovault.org/images/787485
    - https://neurovault.org/images/787490
    - https://neurovault.org/images/787495

#### Motor

- Motor fMRI task paradigm
- Motor sequencing task
- Finger tapping task

Contrast of interest: motor task vs baseline

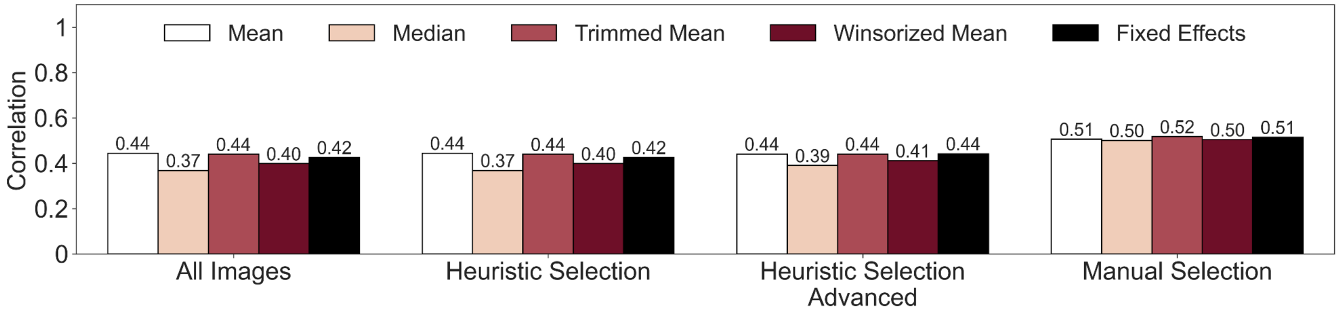


**Figure S5. Evaluation of motor image-based meta-analyses with NeuroVault**. Comparison of the different image selection methods and estimators. The bars represent the correlation with the reference maps.

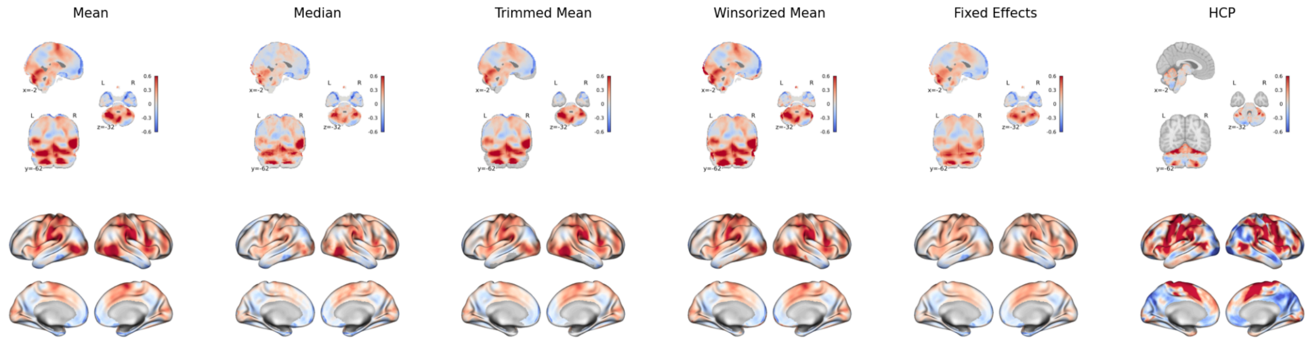


**Figure S6. IBMA result for motor, including all images.**

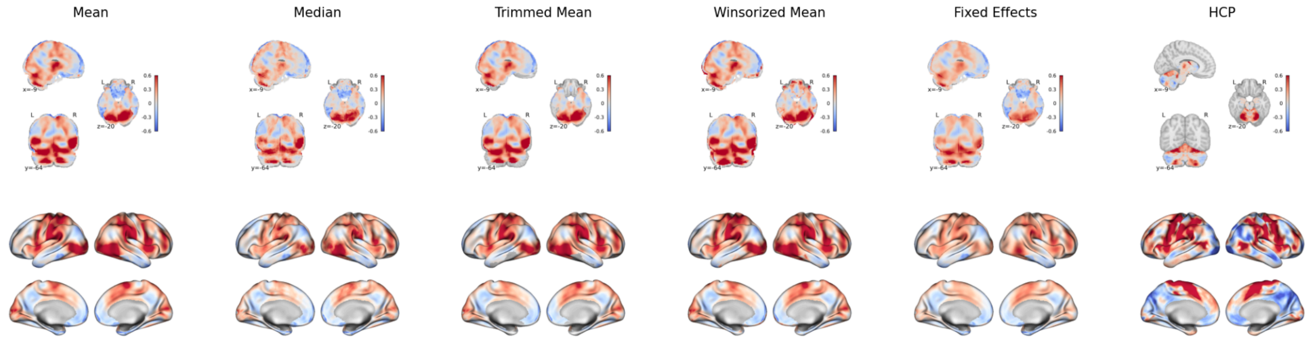


**Figure S7. IBMA result for motor, with automatically selected images.**

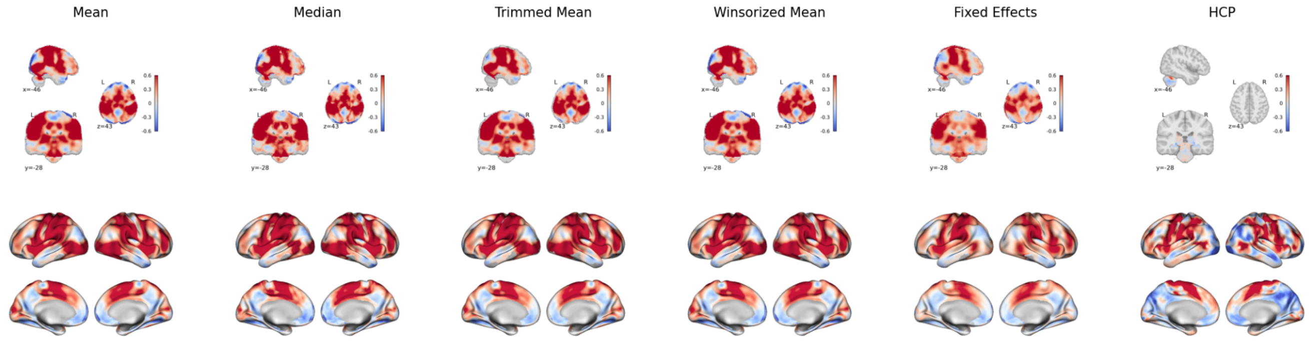


**Figure S8. IBMA result for motor, with manually selected images.**

#### NeuroVault Collections and PMIDs

### 1035

- Link: https://neurovault.org/collections/1035

- Paper: https://pubmed.ncbi.nlm.nih.gov/26635585
- Include: True
- Images:
  - https://neurovault.org/images/14045
    - Selected
  - https://neurovault.org/images/14046
    - Selected

**1904**

- Link: https://neurovault.org/collections/1904
- Paper: https://pubmed.ncbi.nlm.nih.gov/27845254
- Include: True
- Images:
  - https://neurovault.org/images/39070
    - Selected
  - https://neurovault.org/images/39071
  - https://neurovault.org/images/39072
  - https://neurovault.org/images/39073
  - https://neurovault.org/images/39075

**2853**

- Link: https://neurovault.org/collections/2853
- Paper: https://pubmed.ncbi.nlm.nih.gov/28957344
- Include: True
- Images:
  - https://neurovault.org/images/53714
  - https://neurovault.org/images/53715
  - https://neurovault.org/images/53716
  - https://neurovault.org/images/53717
  - https://neurovault.org/images/53718
  - https://neurovault.org/images/53719
  - https://neurovault.org/images/53720
    - Selected
  - https://neurovault.org/images/53721
    - Selected
  - https://neurovault.org/images/53722
    - Selected
  - https://neurovault.org/images/53723
    - Selected

- https://neurovault.org/images/53724
  - Selected
- https://neurovault.org/images/53725
  - Selected
- https://neurovault.org/images/53726
  - Selected
- https://neurovault.org/images/53727
- https://neurovault.org/images/53728
- https://neurovault.org/images/53729
- https://neurovault.org/images/53730
- https://neurovault.org/images/53731
- https://neurovault.org/images/53732
- https://neurovault.org/images/53733
- https://neurovault.org/images/53746
- https://neurovault.org/images/53747
  - Selected
- https://neurovault.org/images/53748
  - Selected
- https://neurovault.org/images/53749
  - Selected
- https://neurovault.org/images/53750
  - Selected
- https://neurovault.org/images/53751
  - Selected
- https://neurovault.org/images/53752
- https://neurovault.org/images/53753
- https://neurovault.org/images/53754
- https://neurovault.org/images/53755
- https://neurovault.org/images/53756
- https://neurovault.org/images/53757
- https://neurovault.org/images/53758
- https://neurovault.org/images/53759
- https://neurovault.org/images/53760
- https://neurovault.org/images/53761
- https://neurovault.org/images/53762
- https://neurovault.org/images/53763
- https://neurovault.org/images/53764
- https://neurovault.org/images/53765
- https://neurovault.org/images/53766
  - Selected
- https://neurovault.org/images/53767
  - Selected

- https://neurovault.org/images/53768
  - Selected
- https://neurovault.org/images/53769
  - Selected
- https://neurovault.org/images/53770
  - Selected
- https://neurovault.org/images/53771
  - Selected
- https://neurovault.org/images/53772
- https://neurovault.org/images/53773
- https://neurovault.org/images/53774
- https://neurovault.org/images/53775
- https://neurovault.org/images/53776
- https://neurovault.org/images/53777
- https://neurovault.org/images/53778
- https://neurovault.org/images/53779
- https://neurovault.org/images/53780
- https://neurovault.org/images/53781
- https://neurovault.org/images/53782
- https://neurovault.org/images/53783
- https://neurovault.org/images/53784
- https://neurovault.org/images/53785

**3117**

- Link: https://neurovault.org/collections/3117
- Paper: https://pubmed.ncbi.nlm.nih.gov/29356166
- Include: True
- Images:
  - https://neurovault.org/images/59400
    - Selected
  - https://neurovault.org/images/59401
  - https://neurovault.org/images/59402

**4208**

- Link: https://neurovault.org/collections/4208
- Paper: https://pubmed.ncbi.nlm.nih.gov/31211363
- Include: True
- Images:
  - https://neurovault.org/images/100643

            - Selected
    - https://neurovault.org/images/100644
        - Selected
    - https://neurovault.org/images/100646
        - Selected
    - https://neurovault.org/images/100647
        - Selected
    - https://neurovault.org/images/68449
    - https://neurovault.org/images/68450
        - Selected
    - https://neurovault.org/images/68451

**6075**

- Link: https://neurovault.org/collections/6075
- Paper: https://pubmed.ncbi.nlm.nih.gov/31797874
- Include: False
    - Eye movement?
- Images:
    - https://neurovault.org/images/305488
    - https://neurovault.org/images/305493
    - https://neurovault.org/images/305496

**9490**

- Link: https://neurovault.org/collections/9490
- Paper: https://pubmed.ncbi.nlm.nih.gov/32219378
- Include: True
- Images:
    - https://neurovault.org/images/442748
        - Selected

**15652**

- Link: https://neurovault.org/collections/15652
- Paper: https://pubmed.ncbi.nlm.nih.gov/37822708
- Include: True
- Images:
    - https://neurovault.org/images/805634
        - Selected
    - https://neurovault.org/images/805635

- Selected
  - [https://neurovault.org/images/805636](https://neurovault.org/images/805636)
  - [https://neurovault.org/images/805637](https://neurovault.org/images/805637)

#### Pain

- Phasic pain stimulation
- Pain monitor/discrimination task

Contrast of interest: high pain vs low pain

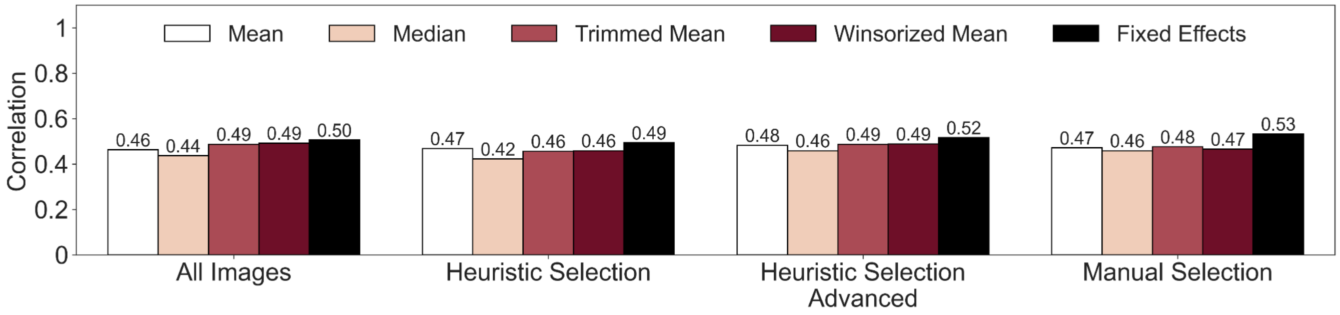


**Figure S9. Evaluation of pain image-based meta-analyses with NeuroVault**. Comparison of the different image selection methods and estimators. The bars represent the correlation with the reference maps.

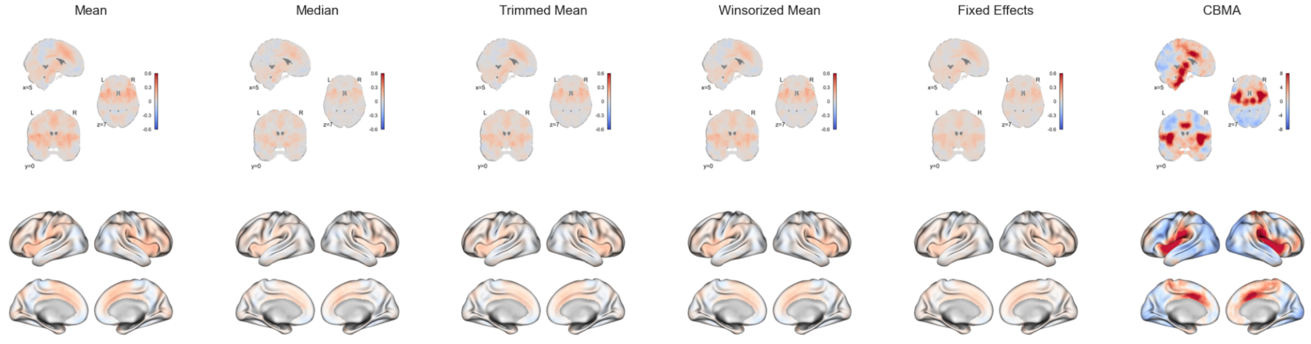


**Figure S10. IBMA result for pain, including all images.**

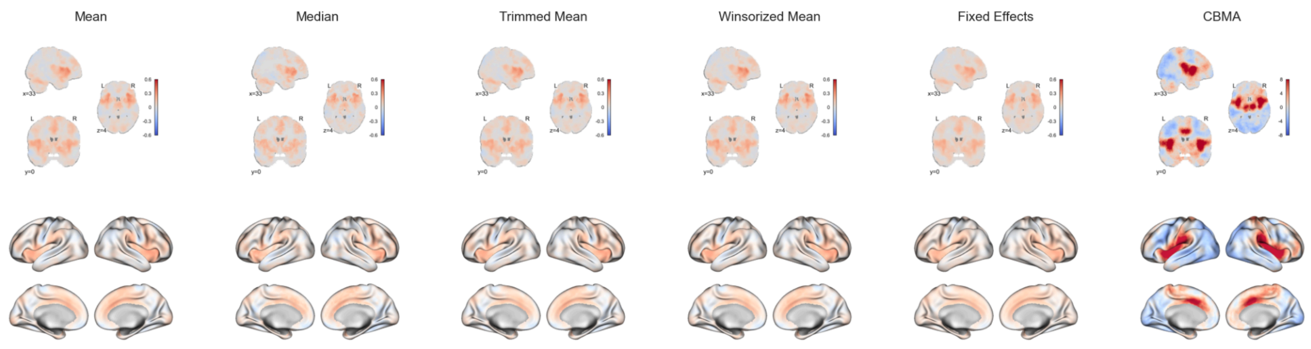


**Figure S11. IBMA result for pain, with automatically selected images.**

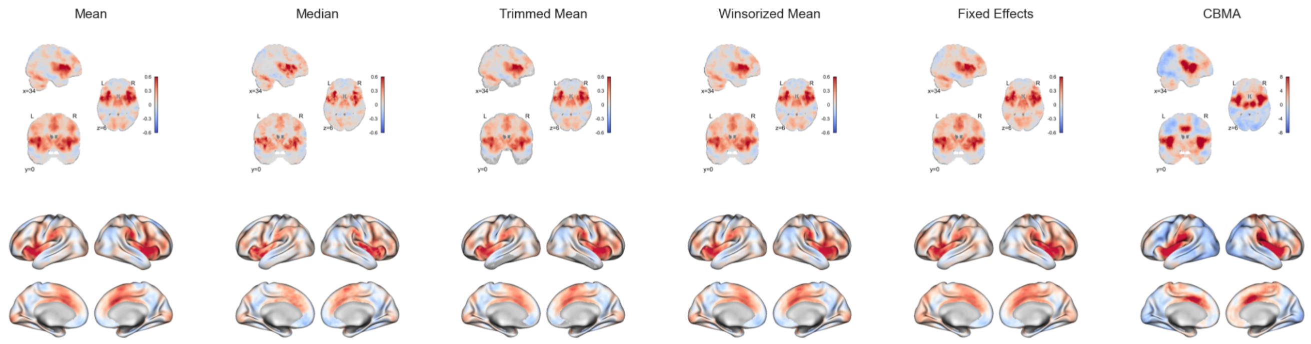


**Figure S12. IBMA result for pain, with manually selected images.**

#### NeuroVault Collections and PMIDs

### 1526

- Link: https://neurovault.org/collections/1526
- Paper: https://pubmed.ncbi.nlm.nih.gov/28852151
- Include: True
    - High vs low thermal pain
- Images:
    - https://neurovault.org/images/29351
        - Selected: pain vs no pain
    - https://neurovault.org/images/29352
        - Selected: pain vs no pain
    - https://neurovault.org/images/29353
        - Selected: pain vs no pain

**2937**

- Link: https://neurovault.org/collections/2937
- Paper: https://pubmed.ncbi.nlm.nih.gov/28991914
- Include: True
- Images:
    - https://neurovault.org/images/53485
        - Selected: pain vs no pain. Check why it includes two session
    - https://neurovault.org/images/53491
    - https://neurovault.org/images/53492

**6545**

- Link: https://neurovault.org/collections/6545
- Paper: https://pubmed.ncbi.nlm.nih.gov/32248233
- Include: False
    - pain empathy task, participants received a cue (2 s) if the electrical shock was directed at themselves. They did a paired t-test.
- Images:
    - https://neurovault.org/images/359732
    - https://neurovault.org/images/359733
    - https://neurovault.org/images/359734
    - https://neurovault.org/images/359735
    - https://neurovault.org/images/359736
    - https://neurovault.org/images/359737
    - https://neurovault.org/images/359752

**9015**

- Link: https://neurovault.org/collections/9015
- Paper: https://pubmed.ncbi.nlm.nih.gov/33186712
- Include: True
    - high temperature stimulation vs low temperature stimulation
- Images:
    - https://neurovault.org/images/426866
        - Selected

**8400**

- Link: https://neurovault.org/collections/8400

- Paper: https://pubmed.ncbi.nlm.nih.gov/33272905
- Include: False
    - The task included images with injuries; no pain was applied during task
- Images:
    - https://neurovault.org/images/394792

**9513**

- Link: https://neurovault.org/collections/9513
- Paper: https://pubmed.ncbi.nlm.nih.gov/33941755
- Include: True
- Images:
    - https://neurovault.org/images/443552
        - Selected

**11088**

- Link: https://neurovault.org/collections/11088
- Paper: https://pubmed.ncbi.nlm.nih.gov/35289367
- Include: True
    - high temperature stimulation vs low temperature stimulation
- Images:
    - https://neurovault.org/images/546329
    - https://neurovault.org/images/548359
        - Selected
    - https://neurovault.org/images/548360
        - Selected
    - https://neurovault.org/images/548366
    - https://neurovault.org/images/548367
    - https://neurovault.org/images/548368
    - https://neurovault.org/images/548369
    - https://neurovault.org/images/548370
    - https://neurovault.org/images/548371
    - https://neurovault.org/images/548372
    - https://neurovault.org/images/548373
        - Selected
    - https://neurovault.org/images/548374
        - Selected
    - https://neurovault.org/images/548375
        - Selected

**10410**

- Link: https://neurovault.org/collections/10410
- Paper: https://pubmed.ncbi.nlm.nih.gov/35658082
- Include: False
- Images:
  - https://neurovault.org/images/510253

**6016**

- Link: https://neurovault.org/collections/6016
- Paper: https://pubmed.ncbi.nlm.nih.gov/35731646
- Include: True
- Images:
  - https://neurovault.org/images/304525
    - Selected: pain vs no pain contrast
  - https://neurovault.org/images/304529
  - https://neurovault.org/images/304530

**12874**

- Link: https://neurovault.org/collections/12874
- Paper: https://pubmed.ncbi.nlm.nih.gov/36317867
- Include: True
  - Thermal pain stimulation
- Images:
  - https://neurovault.org/images/785504
    - Selected: main effect of pain
  - https://neurovault.org/images/785505
  - https://neurovault.org/images/785629
  - https://neurovault.org/images/785630
  - https://neurovault.org/images/785631
  - https://neurovault.org/images/785632

**12827**

- Link: https://neurovault.org/collections/12827
- Paper: https://pubmed.ncbi.nlm.nih.gov/36329014
- Include: False
  - The contrast are not clearly stated in the metadata of the images

- Images:
    - https://neurovault.org/images/785234
    - https://neurovault.org/images/785237
    - https://neurovault.org/images/785240
    - https://neurovault.org/images/785243
    - https://neurovault.org/images/785246
    - https://neurovault.org/images/785249
    - https://neurovault.org/images/785252
    - https://neurovault.org/images/785255

#### Emotion Processing

- Emotion processing fMRI task paradigm

Contrast of interest: negative emotion vs neutral emotion; face vs shape (HCP task developed by Hariri and colleagues (Smith et al. 2007))

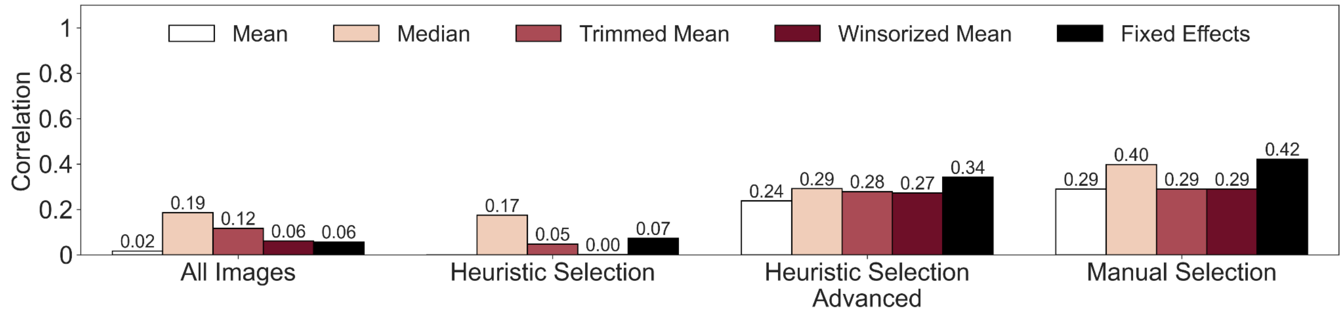


**Figure S13. Evaluation of emotion processing image-based meta-analyses with NeuroVault.** Comparison of the different image selection methods and estimators. The bars represent the correlation with the reference maps.

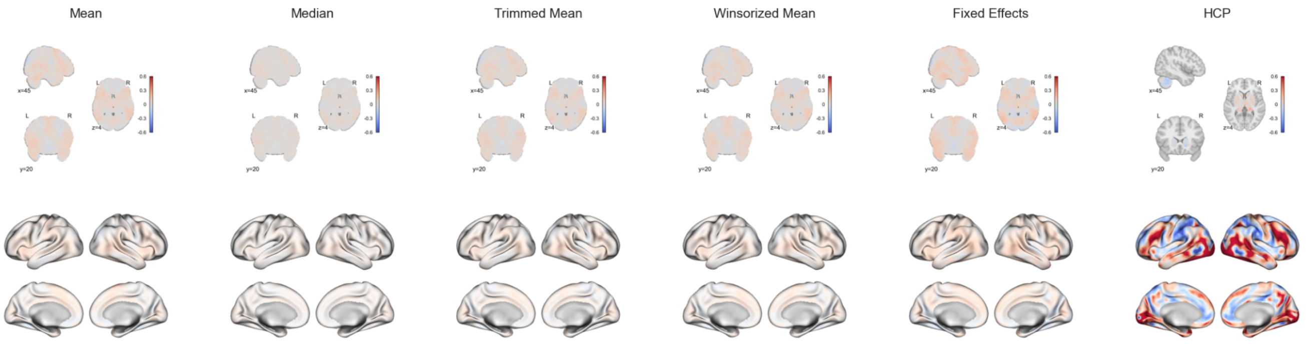

**Figure S14. IBMA result for emotion processing, including all images.**

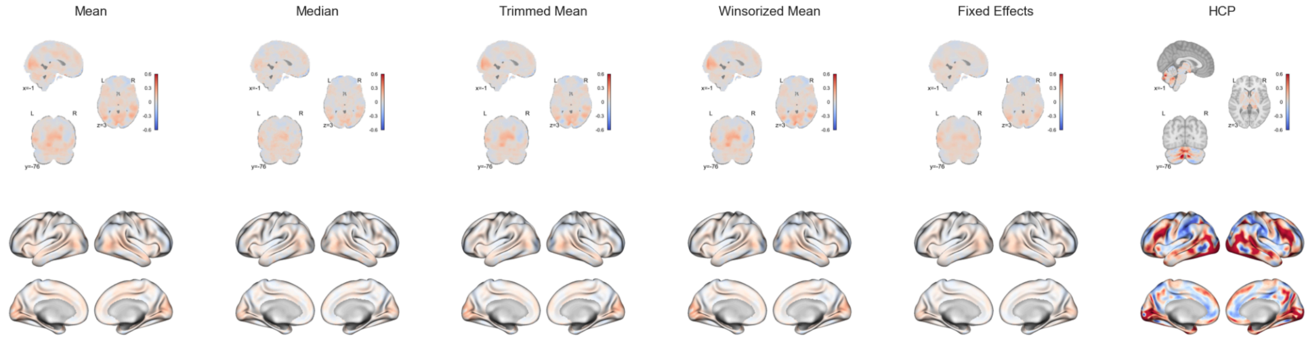


**Figure S15. IBMA result for emotion processing, with automatically selected images.**

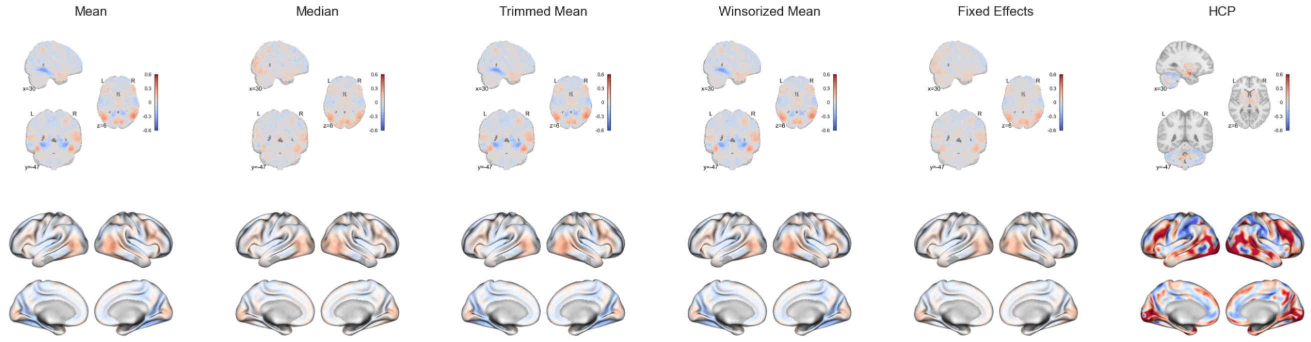


**Figure S16. IBMA result for emotion processing, with manually selected images.**

#### NeuroVault Collections and PMIDs

### 3154

- Link: https://neurovault.org/collections/3154
- Paper: https://pubmed.ncbi.nlm.nih.gov/29447377
- Include: True
  - Positive, negative and neutral faces. In particular they look at positive*negative > neutral
- Images:
  - https://neurovault.org/images/56832
    - Selected: positive*negative > neutral
  - https://neurovault.org/images/56833
    - Selected: positive*negative > neutral
  - https://neurovault.org/images/56834

        - Selected: positive*negative > neutral
    - https://neurovault.org/images/56835
        - Selected: positive*negative > neutral
    - https://neurovault.org/images/56836
        - Selected: positive*negative > neutral
    - https://neurovault.org/images/56837
        - Selected: positive*negative > neutral
    - https://neurovault.org/images/56838
        - Selected: positive*negative > neutral
    - https://neurovault.org/images/56839
    - https://neurovault.org/images/56840
    - https://neurovault.org/images/56841
    - https://neurovault.org/images/56842
    - https://neurovault.org/images/56843
    - https://neurovault.org/images/56844
    - https://neurovault.org/images/56845

**4414**

- Link: https://neurovault.org/collections/4414
- Paper: https://pubmed.ncbi.nlm.nih.gov/30420682
- Include: True
    - Emotional faces paradigm after consuming either alcohol or placebo
- Images:
    - https://neurovault.org/images/100437
        - Selected: faces > houses/shapes
    - https://neurovault.org/images/100438
    - https://neurovault.org/images/100439
    - https://neurovault.org/images/100440

**4096**

- Link: https://neurovault.org/collections/4096
- Paper: https://pubmed.ncbi.nlm.nih.gov/30852995
- Include: False
    - The task is weird it is not a traditional faces > shapes
- Images:
    - https://neurovault.org/images/108829
    - https://neurovault.org/images/108830
    - https://neurovault.org/images/108831
    - https://neurovault.org/images/108832

- https://neurovault.org/images/108833
- https://neurovault.org/images/108835
- https://neurovault.org/images/108836
- https://neurovault.org/images/108837
- https://neurovault.org/images/108838
- https://neurovault.org/images/108839
- https://neurovault.org/images/108876
- https://neurovault.org/images/108877
- https://neurovault.org/images/108878
- https://neurovault.org/images/108879
- https://neurovault.org/images/108884
- https://neurovault.org/images/108885
- https://neurovault.org/images/108886
- https://neurovault.org/images/108887
- https://neurovault.org/images/108888
- https://neurovault.org/images/108889
- https://neurovault.org/images/108890
- https://neurovault.org/images/108891
- https://neurovault.org/images/108892
- https://neurovault.org/images/108893
- https://neurovault.org/images/108894
- https://neurovault.org/images/108895

**6084**

- Link: https://neurovault.org/collections/6084
- Paper: https://pubmed.ncbi.nlm.nih.gov/31812503
- Include: False
    - Not a traditional faces > shapes. Music task
- Images:
    - https://neurovault.org/images/305500
    - https://neurovault.org/images/305501
    - https://neurovault.org/images/305502
    - https://neurovault.org/images/305503

**6237**

- Link: https://neurovault.org/collections/6237
- Paper: https://pubmed.ncbi.nlm.nih.gov/31926280
- Include: False
    - Not a traditional faces > shapes. Movie watching

- Images:
  - https://neurovault.org/images/313222
  - https://neurovault.org/images/313224
  - https://neurovault.org/images/313226
  - https://neurovault.org/images/313228
  - https://neurovault.org/images/313245
  - https://neurovault.org/images/313246
  - https://neurovault.org/images/313256
  - https://neurovault.org/images/313258

**9176**

- Link: https://neurovault.org/collections/9176
- Paper: https://pubmed.ncbi.nlm.nih.gov/33704850
- Include: False
  - This is a meta-analysis
- Images:
  - https://neurovault.org/images/785773

**9246**

- Link: https://neurovault.org/collections/9246
- Paper: https://pubmed.ncbi.nlm.nih.gov/34052365
- Include: False
  - Faces vs shapes, but the contrast look at medication effect and other interactions
- Images:
  - https://neurovault.org/images/440149
  - https://neurovault.org/images/440150
  - https://neurovault.org/images/440151

**11274**

- Link: https://neurovault.org/collections/11274
- Paper: https://pubmed.ncbi.nlm.nih.gov/35657777
- Include: True
  - Threat faces vs shapes
- Images:
  - https://neurovault.org/images/564309
    - Selected
  - https://neurovault.org/images/564310
    - Selected

#### Social Cognition

- theory of mind task: this mostly refers to “functional localizer task for ToM”
- social cognition (theory of mind) fMRI task paradigm: social feedback task

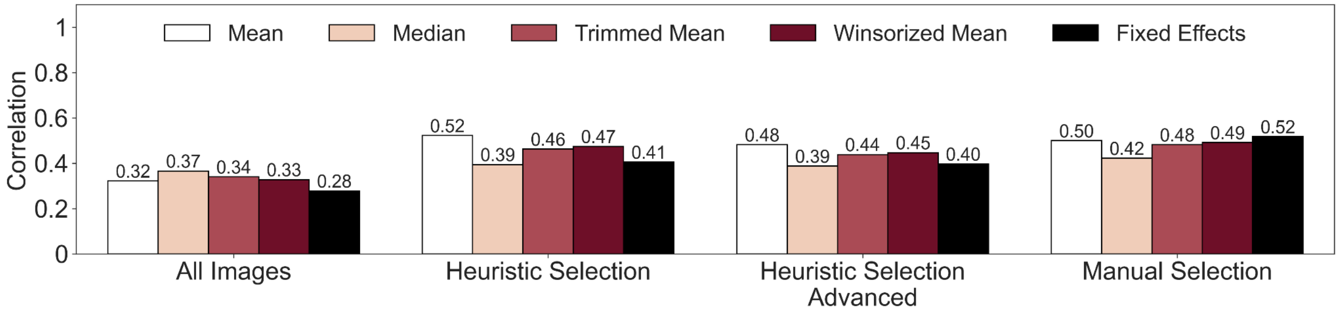


**Figure S17. Evaluation of social cognition image-based meta-analyses with NeuroVault.** Comparison of the different image selection methods and estimators. The bars represent the correlation with the reference maps.

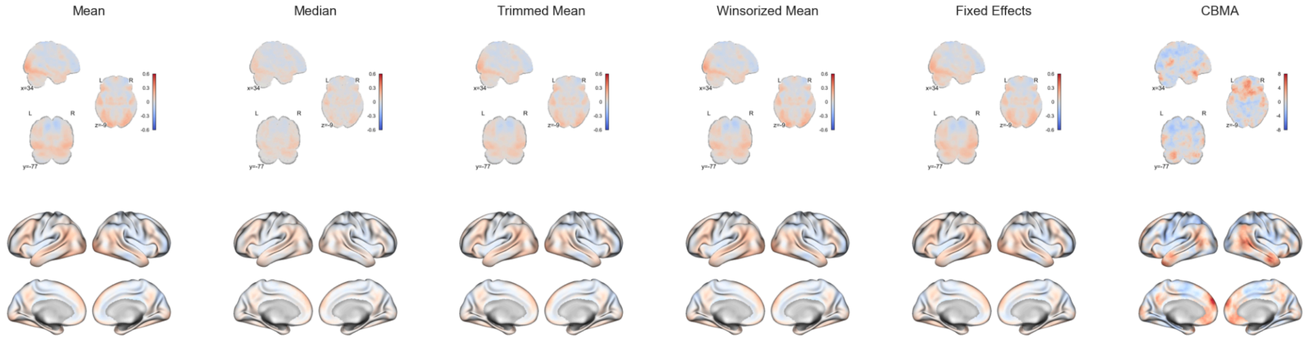


**Figure S18. IBMA result for emotion processing, including all images.**

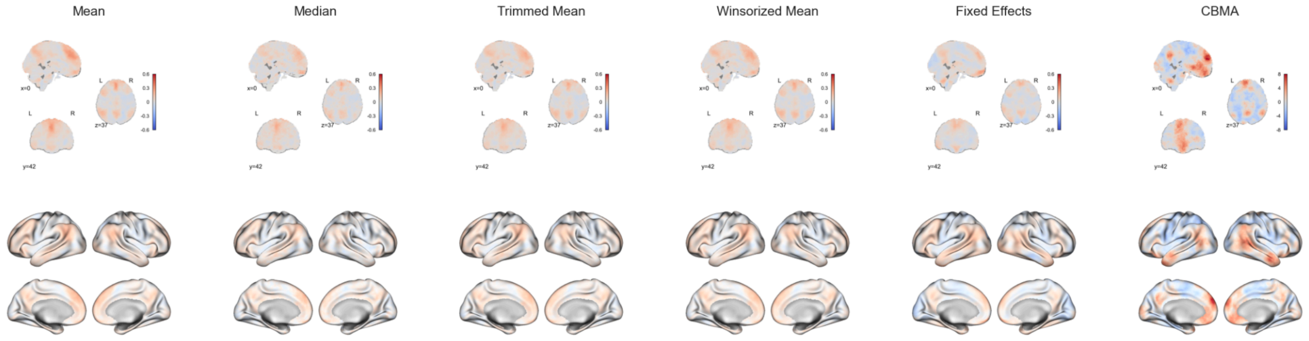

**Figure S19. IBMA result for emotion processing, with automatically selected images.**

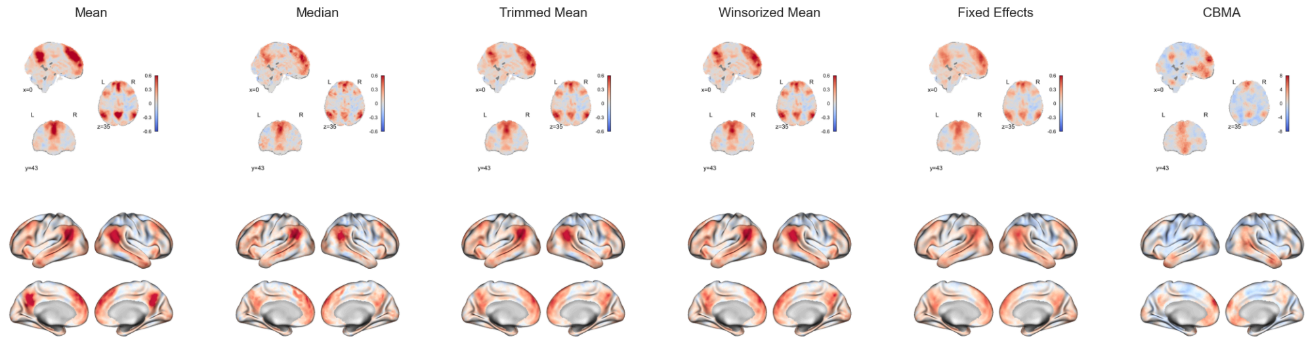


**Figure S20. IBMA result for emotion processing, with manually selected images.**

#### NeuroVault Collections and PMIDs

Note: Look for theory of mind localizer task. TPJ, DMN, etc

### 1377

- Link: https://neurovault.org/collections/1377
- Paper: https://pubmed.ncbi.nlm.nih.gov/22440654
- Include: True
- Images:
    - https://neurovault.org/images/19111
        - Selected: factor belief
    - https://neurovault.org/images/19112
        - Selected: factor desire
    - https://neurovault.org/images/19113

### 1379

- Link: https://neurovault.org/collections/1379
- Paper: https://pubmed.ncbi.nlm.nih.gov/24236763
- Include: True
- Images:
    - https://neurovault.org/images/19122
        - Selected: factor belief
    - https://neurovault.org/images/19123

- Selected: factor desire
  - https://neurovault.org/images/19124
  - https://neurovault.org/images/19125
  - https://neurovault.org/images/19126
  - https://neurovault.org/images/19127
  - https://neurovault.org/images/19128
  - https://neurovault.org/images/19129
  - https://neurovault.org/images/19130
  - https://neurovault.org/images/19131
  - https://neurovault.org/images/19132

**1378**

- Link: https://neurovault.org/collections/1378
- Paper: https://pubmed.ncbi.nlm.nih.gov/25527113
- Include: True
  -
- Images:
  - https://neurovault.org/images/19115
    - Selected: factor of belief
  - https://neurovault.org/images/19116
    - Selected: factor of self
  - https://neurovault.org/images/19117
  - https://neurovault.org/images/19118
  - https://neurovault.org/images/19119
  - https://neurovault.org/images/19120
  - https://neurovault.org/images/19121

**1649**

- Link: https://neurovault.org/collections/1649
- Paper: https://pubmed.ncbi.nlm.nih.gov/26539094
- Include: True
- Images:
  - https://neurovault.org/images/24538
    - Selected: mental interaction

**1478**

- Link: https://neurovault.org/collections/1478
- Paper: https://pubmed.ncbi.nlm.nih.gov/27688764

- Include: False
    - Not a ToM task
- Images:
    - https://neurovault.org/images/23645

**6038**

- Link: https://neurovault.org/collections/6038
- Paper: https://pubmed.ncbi.nlm.nih.gov/29490088
- Include: False
    - Not a ToM task: social feedback task
- Images:
    - https://neurovault.org/images/305035
    - https://neurovault.org/images/305037
    - https://neurovault.org/images/305038
    - https://neurovault.org/images/305041
    - https://neurovault.org/images/305042

**6039**

- Link: https://neurovault.org/collections/6039
- Paper: https://pubmed.ncbi.nlm.nih.gov/30867073
- Include: False
    - Not a ToM task: Social feedback task
- Images:
    - https://neurovault.org/images/305046
    - https://neurovault.org/images/305047
    - https://neurovault.org/images/305048
    - https://neurovault.org/images/305049
    - https://neurovault.org/images/305050
    - https://neurovault.org/images/305051
    - https://neurovault.org/images/305052
    - https://neurovault.org/images/305053
    - https://neurovault.org/images/305054
    - https://neurovault.org/images/305055
    - https://neurovault.org/images/305056
    - https://neurovault.org/images/305057

**7171**

- Link: https://neurovault.org/collections/7171

- Paper: https://pubmed.ncbi.nlm.nih.gov/31680151
- Include: True
  - Theory of mind localizer
- Images:
  - https://neurovault.org/images/381313

**4140**

- Link: https://neurovault.org/collections/4140
- Paper: https://pubmed.ncbi.nlm.nih.gov/31754103
- Include: False
  - Not a ToM task
- Images:
  - https://neurovault.org/images/126761
  - https://neurovault.org/images/126762
  - https://neurovault.org/images/67726
  - https://neurovault.org/images/67728
  - https://neurovault.org/images/67736

**6782**

- Link: https://neurovault.org/collections/6782
- Paper: https://pubmed.ncbi.nlm.nih.gov/32701449
- Include: True
  - social perceptual decision task
- Images:
  - https://neurovault.org/images/371697
    - Selected: group mean, context contrast
  - https://neurovault.org/images/371698
  - https://neurovault.org/images/384210

**6130**

- Link: https://neurovault.org/collections/6130
- Paper: https://pubmed.ncbi.nlm.nih.gov/33666313
- Include: False
  - social cognition task
- Images:
  - https://neurovault.org/images/404974

**9321**

- Link: https://neurovault.org/collections/9321
- Paper: https://pubmed.ncbi.nlm.nih.gov/33746829
- Include: False
  - Not a ToM task: social interaction task
- Images:
  - https://neurovault.org/images/440459

**10920**

- Link: https://neurovault.org/collections/10920
- Paper: https://pubmed.ncbi.nlm.nih.gov/34438254
- Include: True
  - Not a ToM task: Dynamic Inference Task
- Images:
  - https://neurovault.org/images/529711
    - Selected: emotional > non emotional
  - https://neurovault.org/images/529712
  - https://neurovault.org/images/529713

**9352**

- Link: https://neurovault.org/collections/9352
- Paper: https://pubmed.ncbi.nlm.nih.gov/34465915
- Include: True
  - Not a ToM task
- Images:
  - https://neurovault.org/images/440875
    - Selected: mental interaction
  - https://neurovault.org/images/440876
  - https://neurovault.org/images/440877

#### Response Inhibition

- go/no-go task

Note: Looking for inhibition trials (nogo), to examine neural activation supporting changes in effective cognitive control

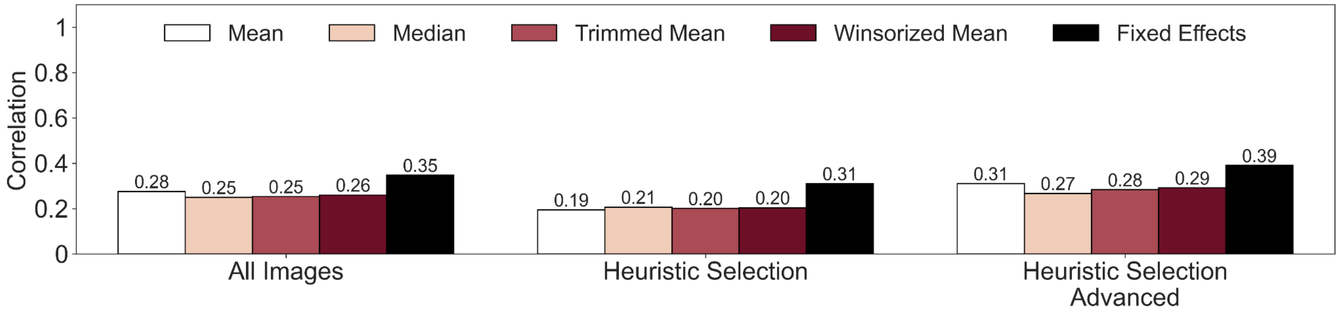


**Figure S21. Evaluation of response inhibition image-based meta-analyses with NeuroVault.** Comparison of the different image selection methods and estimators. The bars represent the correlation with the reference maps.

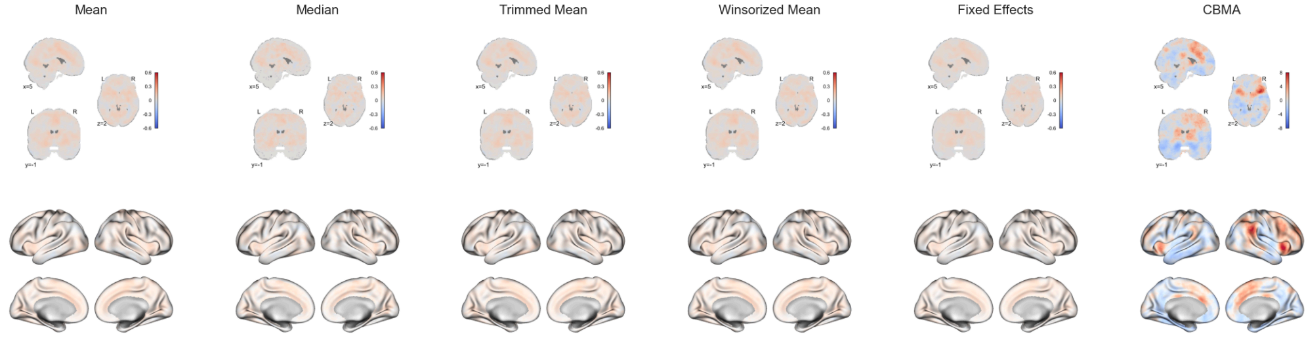


**Figure S22. IBMA result for response inhibition, including all images.**

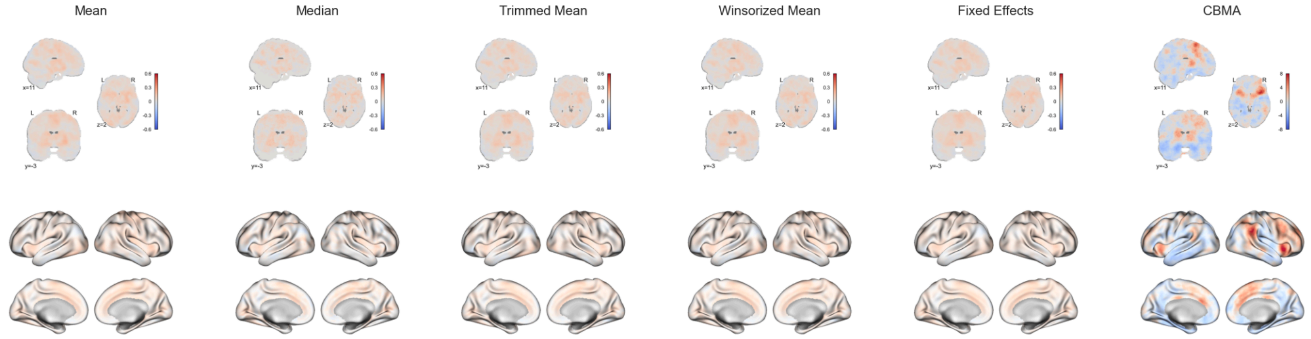


**Figure S23. IBMA result for response inhibition, with automatically selected images.**

#### NeuroVault Collections and PMIDs

### 1827

- Link: [https://neurovault.org/collections/1827](https://neurovault.org/collections/1827)

- Paper: https://pubmed.ncbi.nlm.nih.gov/26434803
- Include: False
    - Use a go/nogo task. Main effect of nogo trials, but they used a longitudinal contrast
- Images:
    - https://neurovault.org/images/28448
    - https://neurovault.org/images/28449
    - https://neurovault.org/images/28450

**1858**

- Link: https://neurovault.org/collections/1858
- Paper: https://pubmed.ncbi.nlm.nih.gov/27445208
- Include: False
    - No nogo trial
- Images:
    - https://neurovault.org/images/28869
        - Go trail
    - https://neurovault.org/images/28870

**2411**

- Link: https://neurovault.org/collections/2411
- Paper: https://pubmed.ncbi.nlm.nih.gov/28392763
- Include: True
    - Nogo trials
- Images:
    - https://neurovault.org/images/43883
        - Note: since this is inhibition the map is negative

**1935**

- Link: https://neurovault.org/collections/1935
- Paper: https://pubmed.ncbi.nlm.nih.gov/28489500
- Include: False
    - Nogo trials, but it tested group differences
- Images:
    - https://neurovault.org/images/29375
    - https://neurovault.org/images/29376
    - https://neurovault.org/images/29377
    - https://neurovault.org/images/29378
    - https://neurovault.org/images/29379

**2698**

- Link: https://neurovault.org/collections/2698
- Paper: https://pubmed.ncbi.nlm.nih.gov/29184096
- Include: True
    - Go/nogo: incentivized cognitive control task in which participants completed go/no-go blocks under high and low financial stakes
- Images:
    - https://neurovault.org/images/51753
    - https://neurovault.org/images/51754
    - https://neurovault.org/images/51755
    - https://neurovault.org/images/51756
    - https://neurovault.org/images/51757
        - Nogo activity: high stake vs low stakes
    - https://neurovault.org/images/53486

**3158**

- Link: https://neurovault.org/collections/3158
- Paper: https://pubmed.ncbi.nlm.nih.gov/29964188
- Include: True
    - Go/nogo task: response to Go/No-Go tasks based on different types of decisions (semantic vs. perceptual) and inputs (words vs. pictures)
- Images:
    - https://neurovault.org/images/57054
        - No Go > Go trials
    - https://neurovault.org/images/57055
    - https://neurovault.org/images/57056
    - https://neurovault.org/images/57057
    - https://neurovault.org/images/57058
    - https://neurovault.org/images/57059
    - https://neurovault.org/images/60655
    - https://neurovault.org/images/60656
    - https://neurovault.org/images/60657
    - https://neurovault.org/images/60659
    - https://neurovault.org/images/63838
    - https://neurovault.org/images/63839
    - https://neurovault.org/images/63840
    - https://neurovault.org/images/63841
    - https://neurovault.org/images/63842

    - https://neurovault.org/images/63843

**4215**

- Link: https://neurovault.org/collections/4215
- Paper: https://pubmed.ncbi.nlm.nih.gov/30957359
- Include: False
    - Go/nogo task: Social Go-NoGo Task
- Images:
    - https://neurovault.org/images/68578
        - Go trials

**5338**

- Link: https://neurovault.org/collections/5338
- Paper: https://pubmed.ncbi.nlm.nih.gov/31396060
- Include: False
    - Go/nogo task: social Go/NoGo cognitive control task. Non nogo trials
- Images:
    - https://neurovault.org/images/127949
    - https://neurovault.org/images/127951
    - https://neurovault.org/images/127955
    - https://neurovault.org/images/129184

**4081**

- Link: https://neurovault.org/collections/4081
- Paper: https://pubmed.ncbi.nlm.nih.gov/31506678
- Include: False
    - Go/No-Go and social Go/No-Go tasks. Non nogo trials
- Images:
    - https://neurovault.org/images/129029
    - https://neurovault.org/images/129030
    - https://neurovault.org/images/65854
    - https://neurovault.org/images/65855
    - https://neurovault.org/images/65856

**5329**

- Link: https://neurovault.org/collections/5329

- Paper: https://pubmed.ncbi.nlm.nih.gov/31747630
- Include: False
    - go/no-go task. Non nogo task
- Images:
    - https://neurovault.org/images/127869
    - https://neurovault.org/images/127870
    - https://neurovault.org/images/127873
    - https://neurovault.org/images/127874

**11190**

- Link: https://neurovault.org/collections/11190
- Paper: https://pubmed.ncbi.nlm.nih.gov/31756586
- Include: True
    - Post-learning Go/NoGo task
- Images:
    - https://neurovault.org/images/550290
        - Maybe: nogo trials
    - https://neurovault.org/images/550291

**5914**

- Link: https://neurovault.org/collections/5914
- Paper: https://pubmed.ncbi.nlm.nih.gov/33316390
- Include: True
    - go/no-go paradigm with stimuli from the conditioning phase
- Images:
    - https://neurovault.org/images/191520
        - Nogo > go
    - https://neurovault.org/images/191521
    - https://neurovault.org/images/191529
        - Nogo > go
    - https://neurovault.org/images/191530

**8086**

- Link: https://neurovault.org/collections/8086
- Paper: https://pubmed.ncbi.nlm.nih.gov/33567443
- Include: True
    - inhibitory control task (Go/NoGo)
- Images:

    - https://neurovault.org/images/392361
        - Nogo > go
    - https://neurovault.org/images/392362
    - https://neurovault.org/images/392363
    - https://neurovault.org/images/392364

**11178**

- Link: https://neurovault.org/collections/11178
- Paper: https://pubmed.ncbi.nlm.nih.gov/34849626
- Include: False
    - motivational Go/NoGo learning task. Nogo contrast not specific
- Images:
    - https://neurovault.org/images/550206
    - https://neurovault.org/images/550207
    - https://neurovault.org/images/550208
    - https://neurovault.org/images/550209
    - https://neurovault.org/images/550210
    - https://neurovault.org/images/550211

**11184**

- Link: https://neurovault.org/collections/11184
- Paper: https://pubmed.ncbi.nlm.nih.gov/38168089
- Include: False
    - motivational Go/ NoGo learning task
- Images:
    - https://neurovault.org/images/550239
    - https://neurovault.org/images/550240
    - https://neurovault.org/images/550241
    - https://neurovault.org/images/550242
    - https://neurovault.org/images/550248
    - https://neurovault.org/images/550249
    - https://neurovault.org/images/563791
    - https://neurovault.org/images/563792
    - https://neurovault.org/images/563793

#### Reward & Decision Making

- Monetary incentive delay
- Gambling task
- Gambling fMRI task paradigm
- Mixed gambles task. Excluding for now, this is mostly collections from the NARPS paper.
- social decision-making task. Excluding

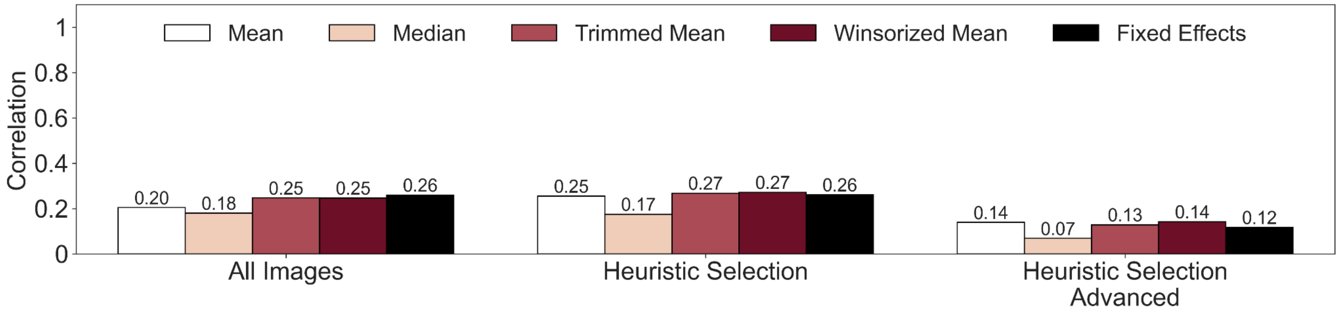


**Figure S24. Evaluation of reward & decision making image-based meta-analyses with NeuroVault.** Comparison of the different image selection methods and estimators. The bars represent the correlation with the reference maps.

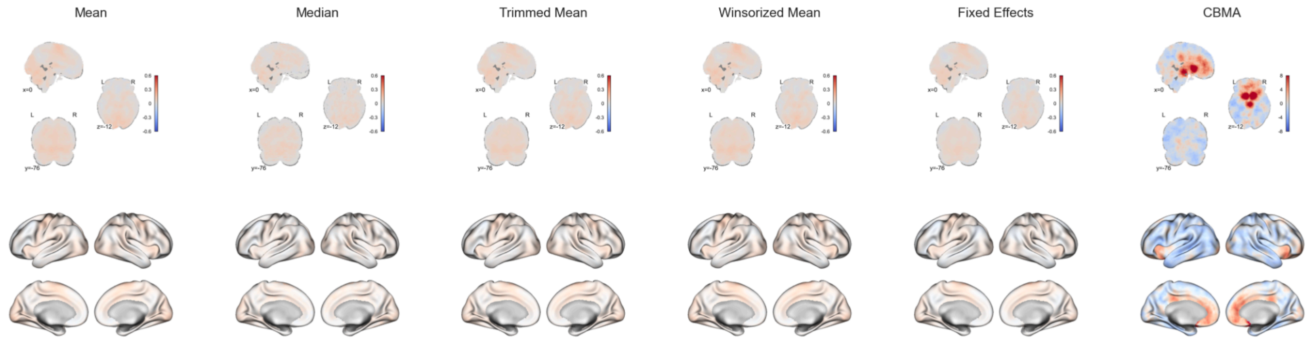


**Figure S25. IBMA result for reward & decision making, including all images.**

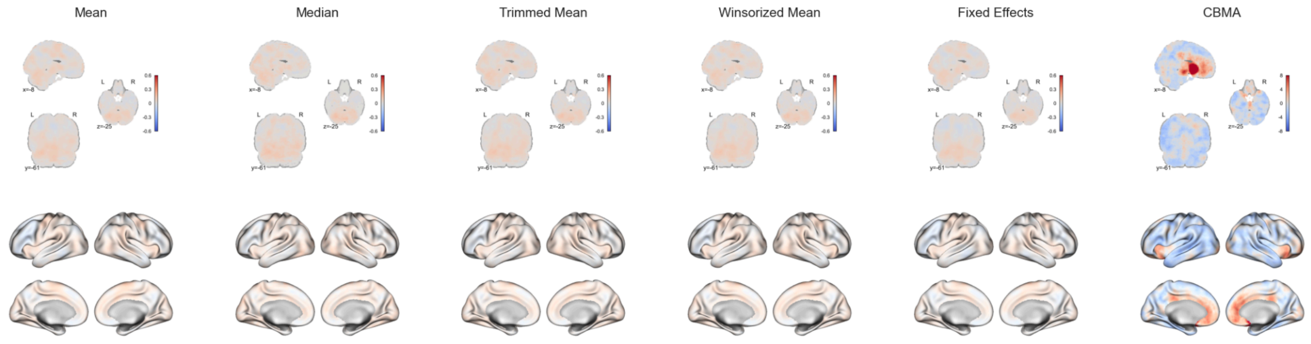


**Figure S26. IBMA result for reward & decision making, with automatically selected images.**

#### NeuroVault Collections and PMIDs

### 4804

- Link: https://neurovault.org/collections/4804
- Paper: https://pubmed.ncbi.nlm.nih.gov/29034316
- Images:
    - https://neurovault.org/images/114245
    - https://neurovault.org/images/114247
    - https://neurovault.org/images/114249
    - https://neurovault.org/images/114334
    - https://neurovault.org/images/114336

### 812

- Link: https://neurovault.org/collections/812
- Paper: https://pubmed.ncbi.nlm.nih.gov/30016481
- Images:
    - https://neurovault.org/images/65082
    - https://neurovault.org/images/65083
    - https://neurovault.org/images/65084

### 6034

- Link: https://neurovault.org/collections/6034
- Paper: https://pubmed.ncbi.nlm.nih.gov/31884223
- Images:
    - https://neurovault.org/images/305019
    - https://neurovault.org/images/305020

- https://neurovault.org/images/305021

**6215**

- Link: https://neurovault.org/collections/6215
- Paper: https://pubmed.ncbi.nlm.nih.gov/32045732
- Images:
  - https://neurovault.org/images/312984
  - https://neurovault.org/images/312985
  - https://neurovault.org/images/312986
  - https://neurovault.org/images/312987

**6282**

- Link: https://neurovault.org/collections/6282
- Paper: https://pubmed.ncbi.nlm.nih.gov/32479377
- Images:
  - https://neurovault.org/images/328379

**5548**

- Link: https://neurovault.org/collections/5548
- Paper: https://pubmed.ncbi.nlm.nih.gov/33030267
- Images:
  - https://neurovault.org/images/129417
  - https://neurovault.org/images/129418

**9057**

- Link: https://neurovault.org/collections/9057
- Paper: https://pubmed.ncbi.nlm.nih.gov/33639845
- Images:
  - https://neurovault.org/images/427285
  - https://neurovault.org/images/427286
  - https://neurovault.org/images/427287
  - https://neurovault.org/images/427288
  - https://neurovault.org/images/427289
  - https://neurovault.org/images/427290
  - https://neurovault.org/images/427291
  - https://neurovault.org/images/427292

- https://neurovault.org/images/427293
- https://neurovault.org/images/427294
- https://neurovault.org/images/427295
- https://neurovault.org/images/427296
- https://neurovault.org/images/427297
- https://neurovault.org/images/427298
- https://neurovault.org/images/427299
- https://neurovault.org/images/427300
- https://neurovault.org/images/427301
- https://neurovault.org/images/427302

**6210**

- Link: https://neurovault.org/collections/6210
- Paper: https://pubmed.ncbi.nlm.nih.gov/33750042
- Images:
  - https://neurovault.org/images/312864
  - https://neurovault.org/images/312865
  - https://neurovault.org/images/312866
  - https://neurovault.org/images/312867
  - https://neurovault.org/images/312868
  - https://neurovault.org/images/312869
  - https://neurovault.org/images/312870
  - https://neurovault.org/images/312871
  - https://neurovault.org/images/312872
  - https://neurovault.org/images/312873
  - https://neurovault.org/images/359857
  - https://neurovault.org/images/359858
  - https://neurovault.org/images/359903
  - https://neurovault.org/images/359904

**8977**

- Link: https://neurovault.org/collections/8977
- Paper: https://pubmed.ncbi.nlm.nih.gov/34263512
- Images:
  - https://neurovault.org/images/408323
  - https://neurovault.org/images/408324
  - https://neurovault.org/images/408325
  - https://neurovault.org/images/408326
  - https://neurovault.org/images/408346

- https://neurovault.org/images/408347
- https://neurovault.org/images/408348
- https://neurovault.org/images/408349
- https://neurovault.org/images/408381
- https://neurovault.org/images/408382
- https://neurovault.org/images/408383
- https://neurovault.org/images/408384
- https://neurovault.org/images/408385
- https://neurovault.org/images/408386
- https://neurovault.org/images/408387
- https://neurovault.org/images/408388
- https://neurovault.org/images/408389
- https://neurovault.org/images/408390
- https://neurovault.org/images/408391
- https://neurovault.org/images/408392
- https://neurovault.org/images/408393
- https://neurovault.org/images/408394
- https://neurovault.org/images/408395
- https://neurovault.org/images/408396
- https://neurovault.org/images/408401
- https://neurovault.org/images/408402
- https://neurovault.org/images/408403
- https://neurovault.org/images/408404
- https://neurovault.org/images/408405
- https://neurovault.org/images/408406
- https://neurovault.org/images/408407
- https://neurovault.org/images/408408
- https://neurovault.org/images/408409
- https://neurovault.org/images/408410
- https://neurovault.org/images/408411
- https://neurovault.org/images/408412
- https://neurovault.org/images/408413
- https://neurovault.org/images/408414
- https://neurovault.org/images/408415
- https://neurovault.org/images/408416

**4199**

- Link: https://neurovault.org/collections/4199
- Paper: https://pubmed.ncbi.nlm.nih.gov/37133998
- Images:

- https://neurovault.org/images/68402
- https://neurovault.org/images/68403

#### Risk

- Balloon analogue risk task

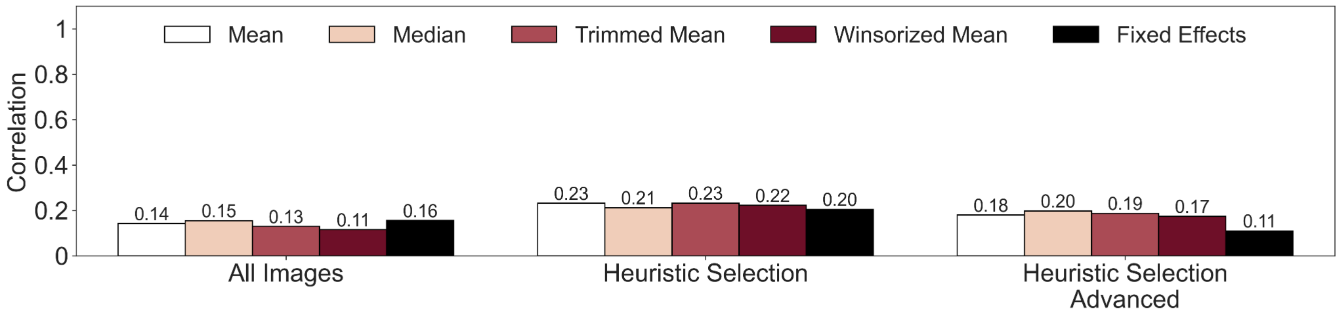


**Figure S27. Evaluation of risk image-based meta-analyses with NeuroVault.** Comparison of the different image selection methods and estimators. The bars represent the correlation with the reference maps.

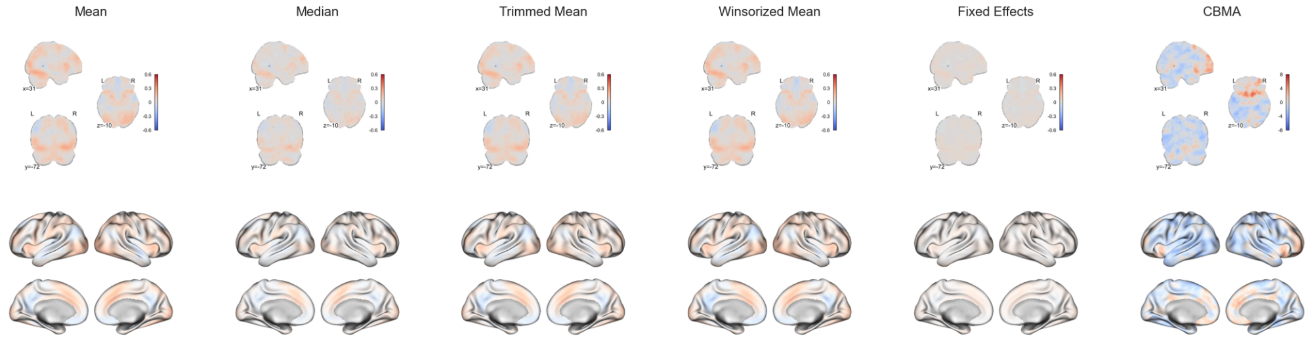


**Figure S28. IBMA result for risk, including all images.**

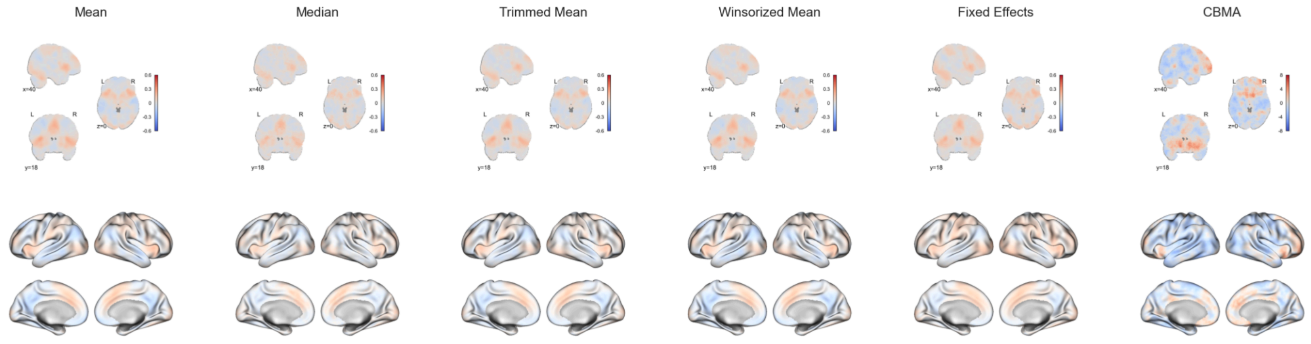


**Figure S29. IBMA result for risk, with automatically selected images.**

#### NeuroVault Collections and PMIDs

### 34

- Link: https://neurovault.org/collections/34
- Paper: https://pubmed.ncbi.nlm.nih.gov/24550270
- Images:
  - https://neurovault.org/images/109

### 1985

- Link: https://neurovault.org/collections/1985
- Paper: https://pubmed.ncbi.nlm.nih.gov/27989774
- Images:
  - https://neurovault.org/images/32001
  - https://neurovault.org/images/32002
  - https://neurovault.org/images/32003
  - https://neurovault.org/images/32004
  - https://neurovault.org/images/32005
  - https://neurovault.org/images/32006
  - https://neurovault.org/images/32007
  - https://neurovault.org/images/32008
  - https://neurovault.org/images/32009
  - https://neurovault.org/images/32010
  - https://neurovault.org/images/32011

### 1873

- Link: https://neurovault.org/collections/1873

- Paper: https://pubmed.ncbi.nlm.nih.gov/28129057
- Images:
    - https://neurovault.org/images/28949
    - https://neurovault.org/images/28950
    - https://neurovault.org/images/28951
    - https://neurovault.org/images/28952

**2606**

- Link: https://neurovault.org/collections/2606
- Paper: https://pubmed.ncbi.nlm.nih.gov/29152222
- Images:
    - https://neurovault.org/images/49974
    - https://neurovault.org/images/49975
    - https://neurovault.org/images/49976
    - https://neurovault.org/images/49977
    - https://neurovault.org/images/49978
    - https://neurovault.org/images/49979
    - https://neurovault.org/images/49980
    - https://neurovault.org/images/49981
    - https://neurovault.org/images/49982
    - https://neurovault.org/images/49983
    - https://neurovault.org/images/49984
    - https://neurovault.org/images/49985
    - https://neurovault.org/images/49986
    - https://neurovault.org/images/49987
    - https://neurovault.org/images/49988
    - https://neurovault.org/images/49989
    - https://neurovault.org/images/49990
    - https://neurovault.org/images/49991
    - https://neurovault.org/images/49992
    - https://neurovault.org/images/49993
    - https://neurovault.org/images/49994
    - https://neurovault.org/images/49995
    - https://neurovault.org/images/49996
    - https://neurovault.org/images/49997

**2039**

- Link: https://neurovault.org/collections/2039
- Paper: https://pubmed.ncbi.nlm.nih.gov/29518712

- Images:
  - https://neurovault.org/images/39173
  - https://neurovault.org/images/39174
  - https://neurovault.org/images/39175
  - https://neurovault.org/images/39176

**3582**

- Link: https://neurovault.org/collections/3582
- Paper: https://pubmed.ncbi.nlm.nih.gov/30575799
- Images:
  - https://neurovault.org/images/62780

**3555**

- Link: https://neurovault.org/collections/3555
- Paper: https://pubmed.ncbi.nlm.nih.gov/30579902
- Images:
  - https://neurovault.org/images/108378
  - https://neurovault.org/images/108393
  - https://neurovault.org/images/108394

**4862**

- Link: https://neurovault.org/collections/4862
- Paper: https://pubmed.ncbi.nlm.nih.gov/31254598
- Images:
  - https://neurovault.org/images/115597
  - https://neurovault.org/images/115598
  - https://neurovault.org/images/115599
  - https://neurovault.org/images/115600

**12312**

- Link: https://neurovault.org/collections/12312
- Paper: https://pubmed.ncbi.nlm.nih.gov/37572094
- Images:
  - https://neurovault.org/images/777557
  - https://neurovault.org/images/777558
  - https://neurovault.org/images/777559

- ○ https://neurovault.org/images/777560

#### Visual Perception

- Passive viewing

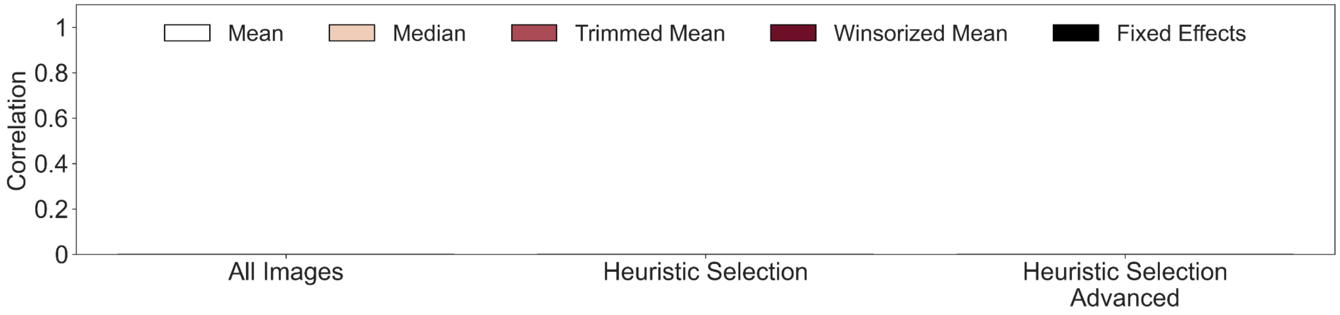


**Figure S30. Evaluation of visual perception image-based meta-analyses with NeuroVault.** Comparison of the different image selection methods and estimators. The bars represent the correlation with the reference maps. Only negative correlations values were found.

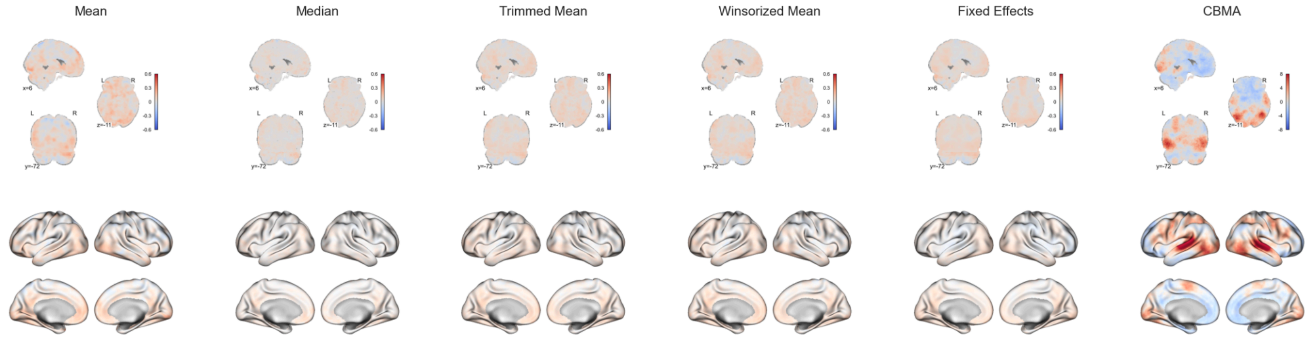


**Figure S31. IBMA result for visual perception, including all images.**

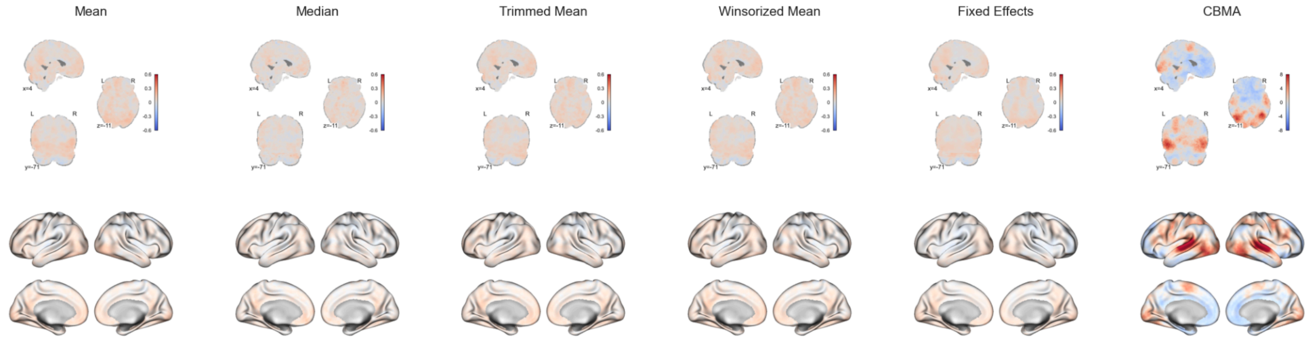


**Figure S32. IBMA result for visual perception, with automatically selected images.**

#### NeuroVault Collections and PMIDs

### 2570

- Link: https://neurovault.org/collections/2570
- Paper: https://pubmed.ncbi.nlm.nih.gov/28803868
- Images:
  - https://neurovault.org/images/49650
  - https://neurovault.org/images/49651

### 2119

- Link: https://neurovault.org/collections/2119
- Paper: https://pubmed.ncbi.nlm.nih.gov/28910669
- Images:
  - https://neurovault.org/images/39882
  - https://neurovault.org/images/39883

### 2715

- Link: https://neurovault.org/collections/2715
- Paper: https://pubmed.ncbi.nlm.nih.gov/30038007
- Images:
  - https://neurovault.org/images/51936
  - https://neurovault.org/images/62435
  - https://neurovault.org/images/65283
  - https://neurovault.org/images/65284
  - https://neurovault.org/images/65285
  - https://neurovault.org/images/65286

- https://neurovault.org/images/65287

**3168**

- Link: https://neurovault.org/collections/3168
- Paper: https://pubmed.ncbi.nlm.nih.gov/30106968
- Images:
  - https://neurovault.org/images/57501

**4096**

- Link: https://neurovault.org/collections/4096
- Paper: https://pubmed.ncbi.nlm.nih.gov/30852995
- Images:
  - https://neurovault.org/images/66992

**3950**

- Link: https://neurovault.org/collections/3950
- Paper: https://pubmed.ncbi.nlm.nih.gov/31408106
- Images:
  - https://neurovault.org/images/112536
  - https://neurovault.org/images/112538
  - https://neurovault.org/images/112539
  - https://neurovault.org/images/112540
  - https://neurovault.org/images/128806
  - https://neurovault.org/images/128808
  - https://neurovault.org/images/65090
  - https://neurovault.org/images/65091
  - https://neurovault.org/images/65092
  - https://neurovault.org/images/65093
  - https://neurovault.org/images/65120

**5182**

- Link: https://neurovault.org/collections/5182
- Paper: https://pubmed.ncbi.nlm.nih.gov/31498505
- Images:
  - https://neurovault.org/images/128002
  - https://neurovault.org/images/128003

- https://neurovault.org/images/128004
- https://neurovault.org/images/128006
- https://neurovault.org/images/128007
- https://neurovault.org/images/128008
- https://neurovault.org/images/128010
- https://neurovault.org/images/128011
- https://neurovault.org/images/128012
- https://neurovault.org/images/128013
- https://neurovault.org/images/128014

**4819**

- Link: https://neurovault.org/collections/4819
- Paper: https://pubmed.ncbi.nlm.nih.gov/33284080
- Images:
  - https://neurovault.org/images/115012
  - https://neurovault.org/images/115015
  - https://neurovault.org/images/115016
  - https://neurovault.org/images/132479
  - https://neurovault.org/images/132480
  - https://neurovault.org/images/132482
